# Mapping specificity, entropy, allosteric changes and substrates in blood proteases by a high-throughput protease screen

**DOI:** 10.1101/2020.07.20.211524

**Authors:** Federico Uliana, Matej Vizovišek, Laura Acquasaliente, Rodolfo Ciuffa, Andrea Fossati, Fabian Frommelt, Matthias Gstaiger, Vincenzo De Filippis, Ulrich auf dem Keller, Ruedi Aebersold

## Abstract

Proteases are among the largest protein families in eukaryotic phylae with more than 500 genetically encoded proteases in humans. By cleaving a wide range of target proteins, proteases are critical regulators of a vast number of biochemical processes including apoptosis and blood coagulation. Over the last 20 years, knowledge of proteases has been drastically expanded by the development of proteomic approaches to identify and quantify proteases and their substrates. In spite of their merits, some of these methods are laborious, not scalable or incompatible with native environments. Consequentially, a large number of proteases remain poorly characterized. Here, we introduce a simple proteomic method to profile protease activity based on isolation of protease products from native lysates using a 96FASP filter and their analysis in a mass spectrometer. The method is significantly faster, cheaper, technically less demanding, easily multiplexed and produces accurate protease fingerprints in near-native conditions. By using the blood cascade proteases as a case study we obtained protease substrate profiles of unprecedented depth that can be reliably used to map specificity, entropy and allosteric changes of the protease and to design fluorescent probes and predict physiological substrates. The native protease characterization method is comparable in performance, but largely exceeds the throughput of current alternatives.

## 1. Introduction

Proteolytic cleavage by proteases is a common protein posttranslational modification and a mechanism that regulates protein functions. It is crucial for cell health and homeostasis and is also involved in the development and progression of different diseases like cancer, inflammation, autoimmune, cardiovascular and metabolic disorders^1, 2^. Therefore, it is not surprising that proteases are widely recognized as diagnostic markers and therapeutic targets in the biomedical field^3^. The knowledge of protease cellular and physiological functions as well as their substrates and cleavage preferences is crucial to design molecules for therapeutic modulation of protease activity^4, 5^. In the wake of the progress achieved by bottom-up mass-spectrometry based proteomics^6^, several techniques to systematically study protease-substrate relationships have been described. They can be grouped into two broad classes. The first aims at concurrently generating activity profiles of numerous proteases present in a complex sample. This is usually accomplished by the use of activity-based probes^7–9^. The second aims at identifying, typically by mass spectrometry, the substrate(s) of specific proteases and this is usually accomplished by exposing a test sample to the protease in question, followed by the analysis of protease products, often referred to as degradomics. Relevant techniques to identify protease substrates include COFRADIC (combined fractional diagonal chromatography)^10, 11^, ChaFraDIC (charge-based fractional diagonal chromatography)^12^, PICS (proteomic identification of protease cleavage sites)^13, 14^ and TAILS (terminal amine isotopic labeling of substrates)^15, 16^ which are reviewed elsewhere^17–19^. More recently, workflows like FPPS (fast profiling of protease specificity)^20^ and especially label-free degradomic workflows like DIPPS (direct in-gel profiling of protease specificity)^21^ and ChaFraTip (ChaFraDIC performed in a pipet tip format)^22^ made protease characterization easier and more accessible by describing simplified workflows and omitting extensive fractionation or labeling steps. Furthermore, DIPPS^21^ and ChaFraTip^22^ can simultaneously map prime and non-prime substrate sites by sequencing the protease-generated peptides. In spite of these developments, several limitations remain which limit the throughput, cost-effectiveness or physiological relevance of these assays. Specifically, the above-mentioned methods meet the following limitations: (i) they assay protease-substrate relationships under less- or non-physiological conditions, e.g. using digested (PICS, ChaFraTip) or denaturated proteins (DIPPS) as substrates; (ii) they require chemical modifications, enrichment or separation steps of the protease products (TAILS, COFRADIC, FPPS, PICS, ChaFraDIC) often resulting in costly, time-consuming and technically demanding protocols; (iii) they are not easily multiplexed and usually limited to capturing a few hundred protease cleavages per experiment/sample/fraction which is often not sufficient to comprehensively cover protease-substrate relationships. The consequences of these limitations are well-reflected in the substrates deposited in the MEROPS protease database, currently the most comprehensive protease-substrate resource^23^. About 60% of the 4000 proteases in MEROPS do not have known substrates (orphan proteases), and less than 200 have more than 30 substrates/cleavages identified to date (Figure 1A) (MEROPS release 12.1). Since coverage of at least 30 substrates/cleavage events is required to calculate a reliable substrate specificity with an error rate of 5%^24^ such analyses are currently only possible for less than 5% of proteases across the kingdoms of life. The expansion of cleavage product datasets generated under near-native conditions improves our understanding of protease-substrate relationships at two levels. First, the large number of substrates will add statistical power to the calculation of reliable protease recognition sequences and highlight the contribution of substrate sterical information to the cleavage pattern. Second, the large number of substrates will aid the training of algorithms to improve the prediction of proteases involved in natural peptide generation, exemplified by Proteasix^25^, PROSPER^26^ and SitePrediction^27^.

**Figure 1.**
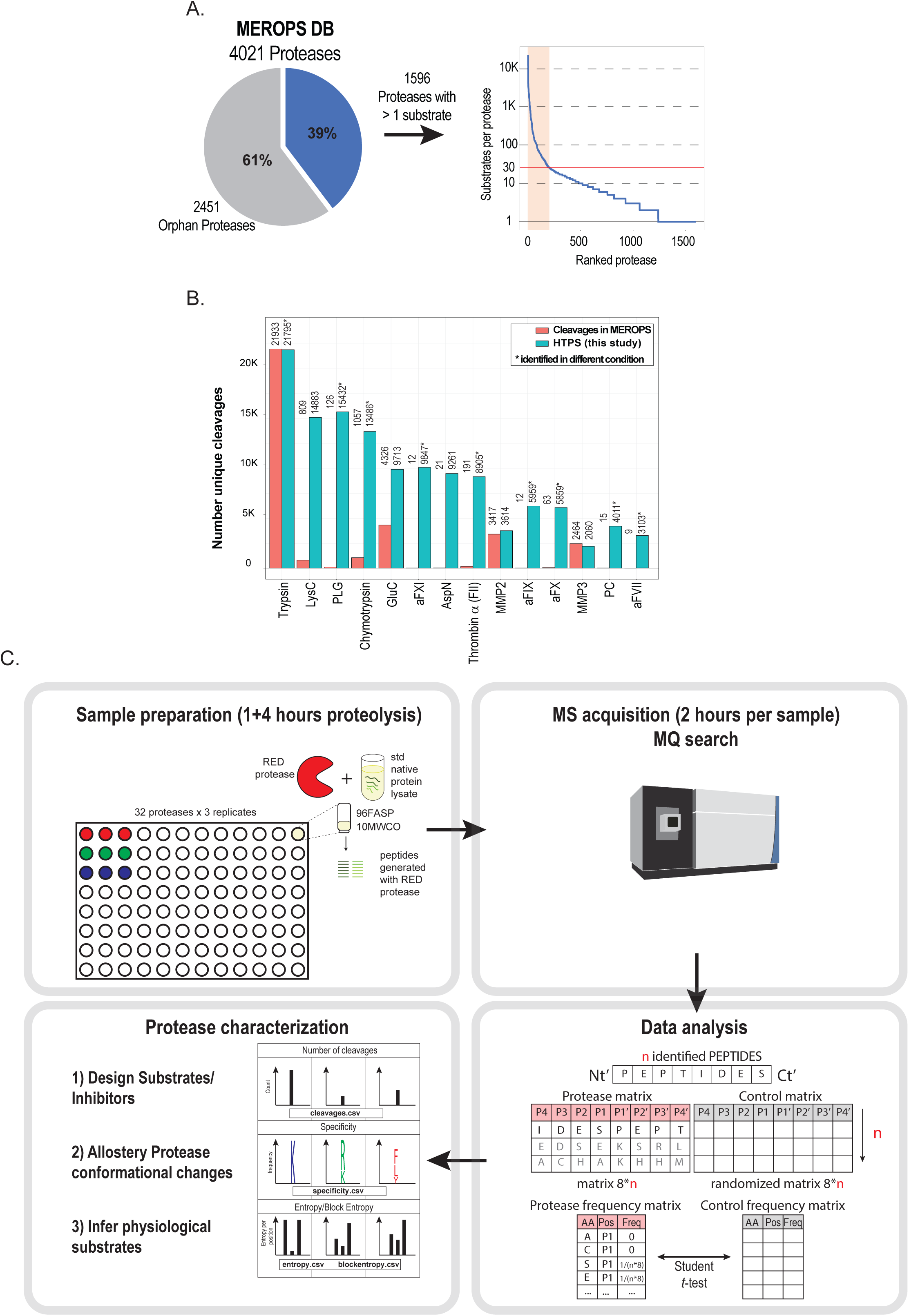
The protease characterization challenge and the HTPS workflow. (A) Distribution of identified substrates per protease annotated in the MEROPS (release 12.1) database. From 4021 proteases reported across the kingdom of life, 2451 are orphan proteases (protease without known substrates) and only around 200 proteases have a sufficient number of known cleavages (i.e. 30 or more) to calculate their specificity with an error rate of 5 % (red line on the chart). (B) Bar plot showing the number of reported protease substrates/cleavages annotated in MEROPS (red) and the number of potential protease substrates/cleavages identified in this study (light green). (C) High-throughput native microscale protease screen (HTPS). In the screen, a native cell lysate is proteolyzed with the studied protease. The protease-generated peptides are collected, analyzed by MS, and the identified substrate peptides are analyzed to retrieve activity, specificity and entropy data. This data can be used to i) design synthetic substrates, ii) characterize allosteric conformational changes and iii) infer physiological protease substrates.

In this study, we developed, benchmarked and applied a streamlined method for high-throughput parallel protease characterization, which we dub “High-Throughput Protease Screen” (HTPS). In contrast to other methods, HTPS is based on the simple isolation of protease-specific peptides from native lysates using a 96 FASP (96 wells filter-aided sample preparation)^28, 29^ that are subsequently identified by data dependent analysis (DDA) mass spectrometry. The method is carried out under near-physiological substrate conditions, obviates labeling and chemical modifications prior or post-protease treatment, fractionation and C18 peptide clean up and, in the current implementation, allows for simultaneous profiling of up to 32 proteases in triplicates. The experimental method is complemented with an optimized data search strategy and deposited R scripts that perform exhaustive protease characterization from identified peptides.

To evaluate the performance of HTPS, we characterized proteases commonly applied in proteomic workflows (Trypsin, Lys-C, Asp-N, Glu-C and Chymotrypsin), as well as the loose-specificity proteases MMP2 and MMP3 and benchmarked these data against other degradomic approaches. We then applied HTPS to identify products of nine blood-activated coagulation proteases (activated α-, β-, and γ-Thrombin, aFVII, aFIX, aFX, aFXI, activated protein C (aPC) and plasmin (PLG); for gene name conversion see Table S1) acting on a native cell lysate. The results expand the repertoire of known substrates/cleavage events for coagulation proteases by about two orders of magnitude, from 428 to 38513 (Figure 1B, Table S2). To further substantiate the potential of HTPS-generated data, we (i) mapped the allosteric effect of Na^+^ on activity, substrate specificity and cleavage entropy of tested blood cascade proteases, with sufficient sensitivity to characterize changes of substrate preference for α-Thrombin in presence or absence of Na^+^; (ii) designed, based on detected specificity features, fluorescent substrates for activated α-Thrombin and aFX and (iii) developed a statistical framework to predict and score potential physiologically relevant substrates starting from HTPS results and applied it to define a large pool of candidate targets of the blood cascade proteases among secreted proteins.

Hence, we describe a simple, high throughput experimental and computational method for protease product profiling under physiological conditions. We show that the method is robust and sensitive enough to support the data-driven reconstruction of protease recognition sequences, substrate design, prediction of protease substrates and an assessment of the effects of allosteric changes on substrate specificity.

## 2. Results

### 2.1. A method for high-throughput screening of protease substrates and cleavage sites on native proteins

The high-throughput protease screen (HTPS) protocol consists of two main steps: i) sample preparation and data acquisition, ii) computational identification and analysis of protease cleavage products (Figure 1C). First, a native protein lysate is prepared, where endogenous proteases are blocked with low-molecular weight inhibitors and the excess inhibitors as well as peptides resulting from background proteolysis are removed using membrane filters with a 10kDa MWCO. To screen the protease of interest under microscale conditions, 50 µg aliquots of the thus prepared native lysate are proteolyzed with the protease in question at 1:50 enzyme to substrate ratio. This step is performed in 96FASP filter plates with a MWCO of 10 kDa^29^, which retains undigested proteins and the added protease and supports recovery of the cleavage products in the flow-through. Four downstream sample processing steps typical for bottom-up proteomics, namely reduction and alkylation, trypsin digestion and C18 cleanup are bypassed. The procedure preserves native substrate fold and disulfide bridges as these can impact substrate accessibility while performing proteolysis in near-native conditions. The generated samples are free from detergent and salt and the peptides collected after FASP centrifugation are directly analyzed by DDA-MS. This simplifies the sample preparation and the workflow eliminates steps that can lead to peptide loss, induce bias towards a particular class of peptides or alter the protease fingerprints. While trypsinization could be potentially beneficial in a double step proteolysis, we found that the investigated proteases generated a considerable number of peptides and only a generally minor, although variable, amount of under-digested peptides/protein fragments was detected at the end of the reverse phase chromatograms. This does not generally have an impact on chromatography or MS instrument performance, as assessed by the stability of the retention time and relative stability of MS1 intensity of the external standard peptide [Glu1]-fibrinopeptide B (Figure S1A, S1B). The net result of these steps are sets of fragment ion spectra of peptides that are highly enriched for substrates of the protease tested.

In the second step, the protease-generated peptides can be identified with any tandem mass spectra search tool. In our implementation, we used the Andromeda^30^ search engine in MaxQuant^31^ using unspecific database search parameters as described elsewhere^21^. Importantly, we searched the data with a reduced database (*HTPS_DB.fasta*) generated from proteins identified in Trypsin, Lys-C, Asp-N, Glu-C and Chymotrypsin samples. The rationale for this strategy is that searches against large databases with many proteins that are not present/detected in the samples lead to a FDR inflation and a decrease in the PSM^32, 33^, particularly pronounced in case of unspecific searches. In our benchmark, we used *HTPS_DB.fasta* containing 2557 protein sequences corresponding to ∼12 % of the human Uniprot database showing the same distribution of amino acids (Figure S2A, S2B) as the whole Uniprot database. With a FDR control at peptide level set to 0.01, the use of *HTPS_DB.fasta* increased the number of PSMs compared to full a proteome database by 19% in the case of Trypsin and by more than 33% in the case of Chymotrypsin (Figure S2C). This increased the ratio of matched MS/MS spectra over all MS/MS spectra and the number of identified peptides (Figure S2D), while decreasing analysis time (Table S3). Next, positional frequency of amino acids, the cleavage entropy^34^ (a quantitative measure of protease specificity) and block entropy^35^ (a measure of protease sub-site cooperativity) were calculated with a series of scripts that we developed for the study and that were extensively annotated and deposited in GitHub (https://github.com/anfoss/HTPS_workflow). Briefly, after filtering the MQ peptide results (contaminants, decoys and low-score peptides), the cleavage sequences of the identified peptides were converted to a frequency matrix covering 8 amino acids upstream and 8 downstream the cleavage site (P8-P1 and P1’-P8’, respectively). Protease specificity was inferred by comparing the observed frequency with a random (null) distribution generated from the database and computing a two-side paired *t*-test. The cleavage sequences from peptides identified after proteolysis were directly used as input for the protease characterization, as we did not introduce a bias from the original protein termini (Figure S3A-D). Overall, the combined experimental and data analysis HTPS workflow supports the identification of thousands of cleavage events, outperforming, for almost all proteases, the number of reported substrates/cleavages in MEROPS database. This is particularly noteworthy in the case of aFXI, aFIX, aFX, aPC and aFVII proteases for which, so far, less than 30 substrates/cleavages were identified (Figure 1C, Table S2).

### 2.2. Benchmarking the performance of the HTPS screen

To test the performance of HTPS we conducted two distinct benchmarking experiments. First, we applied our protocol to the proteases Trypsin, Lys-C, Asp-N, Glu-C and Chymotrypsin which are specific, well characterized^36^ and commonly used in proteomic workflows. Protease characterization was performed in triplicates using a [E]/[S] ratio of 1:50 on the native lysate as substrate sample. Using the workflow described above we identified a higher number of cleavage events for proteases specific for basic amino acids compared to proteases with other cleavage specificities: we identified around 16600 and 14000 cleavages with trypsin and Lys-C, respectively, with an overlap between the triplicates of 91.6 % and 86.8 %. (Figure 2A). For proteases recognizing amino acids with acidic side chains (Glu-C and Asp-N) and for proteases with lower specificity like Chymotrypsin, we recovered between 8800 and 9400 cleavages, with similar levels of reproducibility (average overlap of 88%) between triplicates (Figure 2A). Non-tryptic peptides often have worse chromatographic separation and fragmentation properties than trypsin products and it is estimated that only 4% of all proteomic data sets are generated with proteases other than trypsin^37^. Nevertheless, the fraction of matched MS/MS spectra over all MS/MS spectra range between 14% - 27% for all analysed proteases (Table S3). From the list of identified cleavages we generated specificity profiles *via* iceLogos^38^ with a *p*-value cutoff of 0.01 (Figure 2B, Table S4, S5 for heat maps see Figure S4) in agreement with data from the MEROPS database^23^ and in-line with their well characterized cleavage specificity profiles^36^.

**Figure 2.**
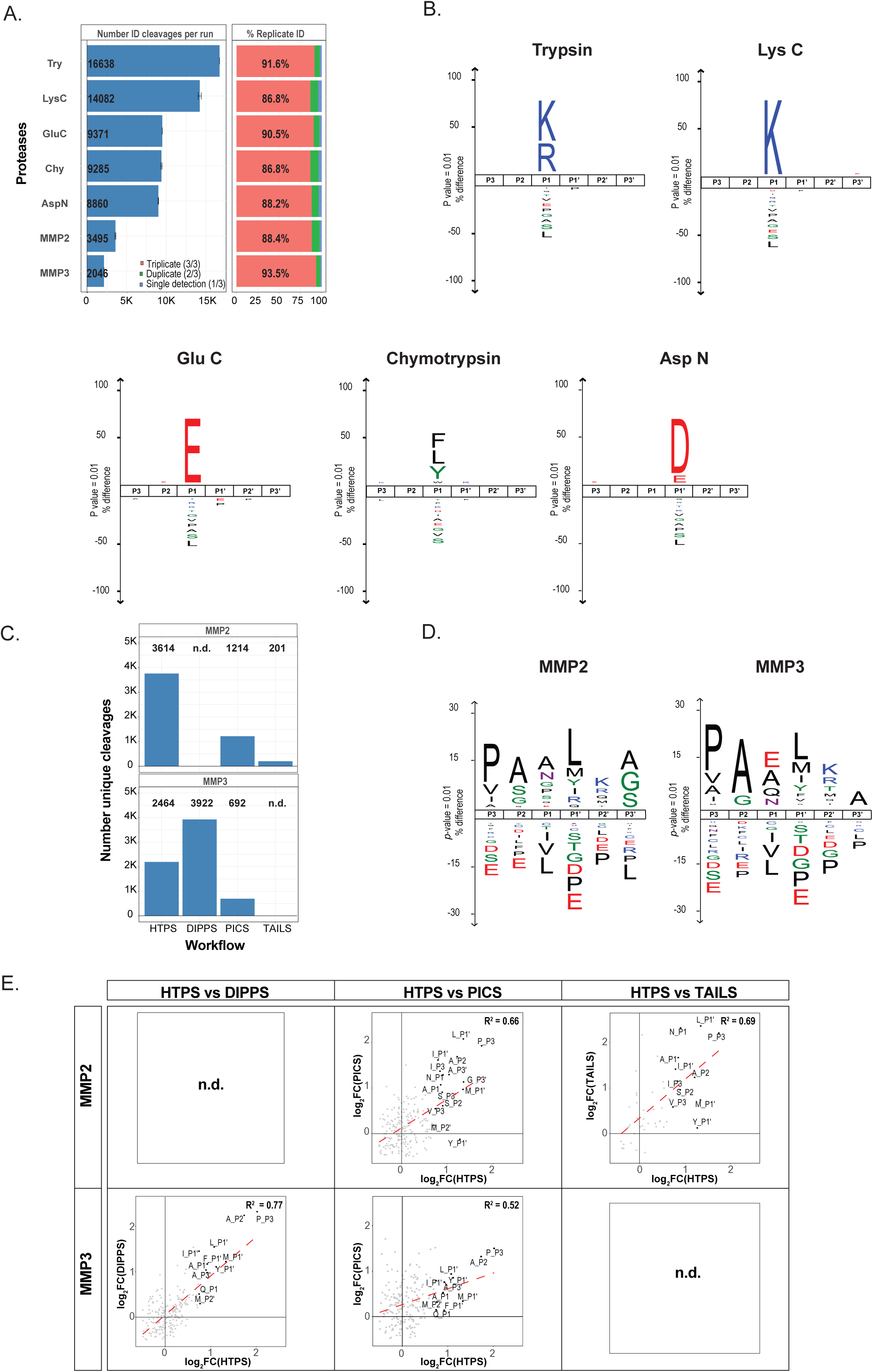
Benchmark of HTPS performance with different proteases. (A) Cleavages generated from benchmark measurements using well-characterized proteases; each protease is characterized by the average number of cleavages identified from three independent replicate experiments (left) and by the overlap across triplicates (right). (B) Specificity benchmark with proteases commonly used in proteomics workflows presented as iceLogos. The protease specificity preferences are shown for P3-P3’ positions. (C) Identified cleavages for MMP2 and MMP3 using different protease characterization approaches (PICS^41, 42^, TAILS^43^, DIPPS^21^ and HTPS). (D) MMP2, MMP3 substrate specificity presented as iceLogos covering P3-P3’ positions. (E) Correlation of the reported specificity enrichment per position for MMP2 and MMP3 between HTPS and other protease workflows (PICS^41, 42^, TAILS^43^, DIPPS^21^).

Second, to further benchmark the performance of HTPS against established methods, we used it to characterize the substrate specificity of the matrix metalloproteases (MMPs) MMP2 and MMP3. MMPs are an important group of proteases that have been extensively studied by proteomic techniques^39^ because of their involvement in development and progression of different pathologies, especially cancer^40^. Over the last few years, the substrate specificity of MMPs was characterized using different methods, including PICS^41, 42^, TAILS^43^ and DIPPS^21^. For the comparison, we used the cleavages identified in the studies and analyzed them with the HTPS workflow to generate the frequency matrices as the basis for the respective positional specificities. In three out of four comparisons our approach resulted in a higher number of cleavages (Figure 2C), 3614 for MMP2 and 2464 for MMP3 and an overlap of almost 90% between triplicates (Figure 2A, Table S4). We determined all positional AA enrichments in comparison to the natural AA distribution in the database, reporting only significant values (adjusted *p-*value < 0.01, Figure 2D, Table S5). HTPS data indicated that MMP2 and MMP3 both have similar specificities (Figure 2D), with preference for Pro, Ala, Val and Ile at P3 position. Further, at P2 position, we observed a preference for Ala, Ser, Gly for MMP2 and Ala, Gly for MMP3. At P1 position Ala, Asn and Pro was observed for MMP2 and Glu, Ala, Gln and Asn for MMP3. Additionally, we observed a preference for Leu, Met, Tyr, Ile at P1’ and Lys, Arg, Met, Thr at P2’ for both proteases and a different specificity in position P3’ for Ala (in case of MMP3) and for Ala, Gly and Ser (in case of MMP2), which is mostly in agreement with the proteases preference reported in other aforementioned studies^21, 41–43^. The comparison of the methods in terms of reported positional amino acid enrichments showed a good overall correlation between PICS, TAILS, DIPPS and HTPS with the R^2^ ranging from 0.52 for PICS to 0.77 for DIPPS (Figure 2E). Taken together, these observations corroborate the validity of HTPS as an alternative method for protease profiling.

### 2.3. High-throughput screening of blood coagulation cascade proteases

To fully exploit the potential of the method, we applied it to comprehensively characterize the blood cascade serine proteases. The group of enzymes tested consists of blood coagulation proteases aFVII, aFIX, aFX, aFXI, activated α-Thrombin, PLG and aPC as well as Thrombin isoforms β-, γ-, that are autoproteoytic products of α-Thrombin. We chose these proteases because they (i) are biologically and chemically related; (ii) have a substantial therapeutic potential; (iii) have been to some extent structurally characterized and (iv) their repertoire of substrates is not yet fully characterized. The concerted action of these serine proteases regulates blood clot formation through activation of Thrombin which converts fibrinogen to insoluble fibrin and activates platelets *via* PAR1 proteolytic activation^44^. Besides the nine blood coagulation proteases we also included chymotrypsin to the screen because it has the archetypal protease structure for the S1 chymotrypsin-like family^45^.

The respective proteases were analyzed using the HTPS workflow. The detected specificity features are summarized in Figure 3A. For each protease we identified cleavages ranging from 1800 for aFVII and up to more than 10000 for PLG (Figure 3B). This represents an increase in the number of identified cleavages, by about two orders of magnitude compared to MEROPS database (428 vs 38513 unique substrates/cleavages, Figure 1C, Table S2). While most of these substrates are not likely to be processed during blood coagulation due to the nature of the substrate sample, they are nevertheless very useful to determine the cleavage specificity, entropy, allostery and other functional/structural properties of the proteases. The heat maps shown in Figure 3A and Figure S4 report significant (corrected *p*-value<0.01, Table S5) enrichment of amino acids around the cleavage site for positions P4-P4’, in comparison to the amino acid distribution in *HTPS_DB*. To gain a more structured insight into the cleavage specificity relationships among the tested proteases, we performed an unsupervised hierarchical clustering according to their substrate preferences (Figure 3C). This analysis revealed the existence of 4 clusters. Cluster 1 included PLG, aFXI and γ-Thrombin, proteases with a strict specificity limited to position P1 for Arg and Lys. Cluster 2 included α- and β-Thrombin, and aFVII and showed specificity in P1 for Arg and a contribution to the specificity of all positions close to the cleavage site (P3-P2’). Cluster 3 contained aFX, aPC and aFIX and showed an intermediate specificity between the first two clusters, but generally closer to cluster 1. As expected, chymotrypsin clustered separately from the clotting proteases as it shows a cleavage specificity for hydrophobic amino acids (Phe, Trp and Tyr at P1 position and Leu, Met to a lesser extent).

**Figure 3.**
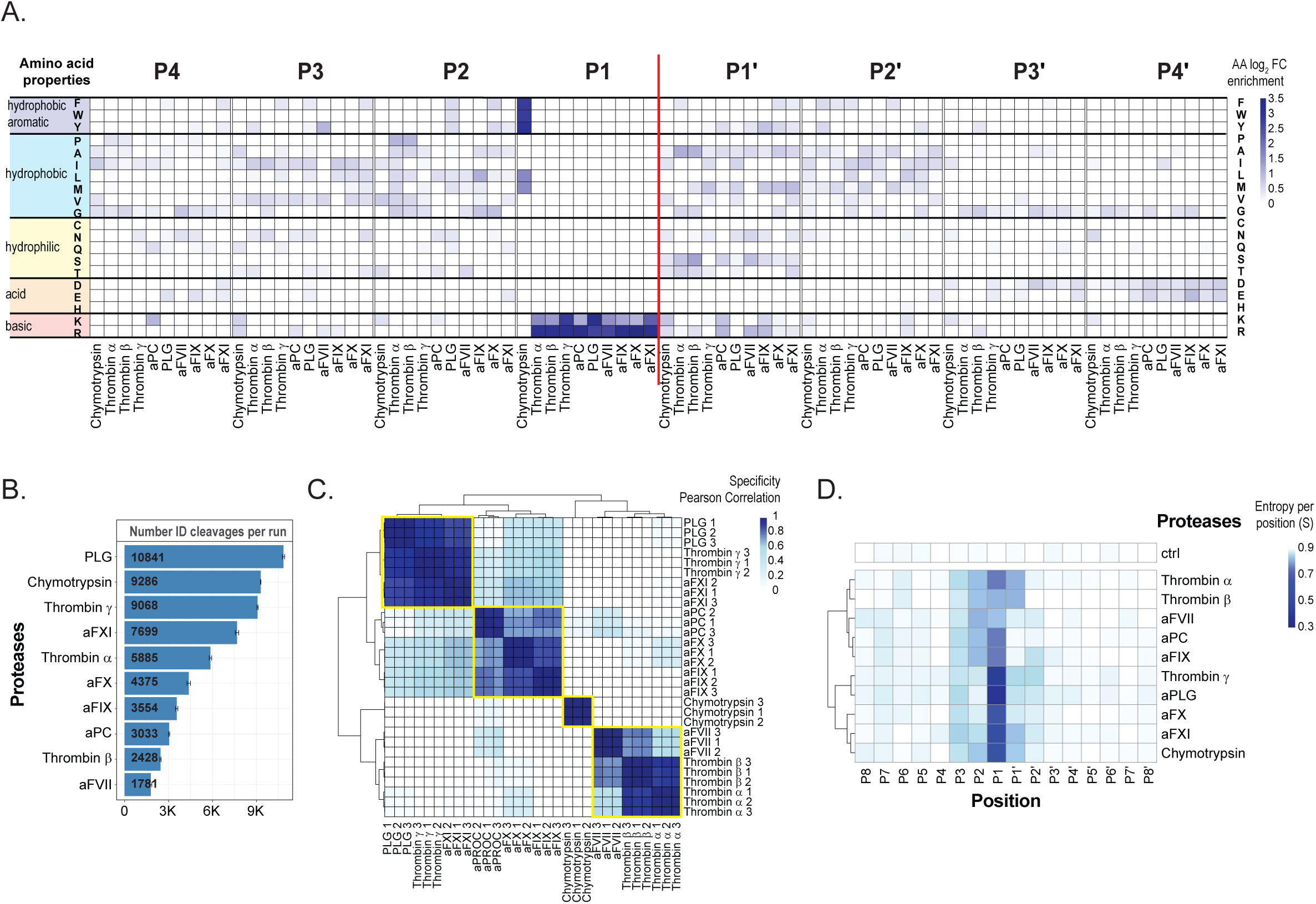
High-throughput screening of coagulation proteases. (A) Positional substrate preferences of coagulation protease from the chymotrypsin-like family proteases (Chymotrypsin, activated α-, β-, γ-Thrombin, aFVII, aFIX, aFX, aFXI, aPC and PLG). The heat map includes positions P4-P4’. The AAs are grouped according to their physicochemical properties. The enrichments are reported as log_2_ FC compared to a random AA distribution generated from HTPS database. (B) Cleavages generated from coagulation proteases; each protease is characterized by the average number of cleavages identified from three independent replicate experiments. (C) Unsupervised hierarchical cluster of coagulation proteases. The color scale describes the Pearson correlation coefficient value calculated for the respective protease samples. (D) Unsupervised hierarchical cluster of coagulation proteases according to the positional cleavage entropy. The color scale describes the entropy (S) values for positions P8-P8’ for all respective proteases included in the assay.

Both, specificity profiles and clustering are in-line with the prior knowledge about these proteases. All coagulation proteases have a defined trypsin-like specificity in position P1. There is also a strong preference for substrates with Arg and to a lesser extent for Lys in P1 position where specificity is tightly regulated by the ionic interaction between the negative carboxylate group of Asp-189 and the positive charged group of the substrate^46, 47^. While the detected enrichment of Arg at P1 position was similar for all proteases, the level of enrichment of Lys was highest for the members of cluster 1. All profiles are characterized by higher specificity in P1 position and lower specificity in other extended positions, indicating that, similar to Trypsin and Chymotrypsin, the protease specificity is determined mainly by the P1 position. This is also evident from the sub-pocket resolved cleavage entropy profiles (Figure 3D), which show the substrate preference per position for each protease. Proteases from cluster 1 are promiscuous proteases, their specificity is essentially determined by the amino acid in position P1 and, as consequence, they cleave more frequently compared to other coagulation proteases (Figure 3B).

Coagulation proteases differ from chymotrypsin structurally by the presence of two insertion loops (loop 60 and loop 148)^44^. These loops form a rigid lid-like structure which regulates accessibility to the catalytic pocket, generates a more extended specificity for coagulation proteases (from P3 to P2’) and thus a distinct substrate fingerprint. As an example, the preference of aFX for Gly in position P2 (Figure 3A) is generated by bulky residues in the insertion loop which accepts small amino acids at the corresponding substrate positions^48^. In α-Thrombin, the 60-loop generates the preference in position P2 for Pro, hydrophobic and planar residues and a preference in position P1’ for small residues like Ala (Figure 3A). The key role of the steric hindrance of the 60-loop in the selectivity of α-Thrombin was shown by mutagenesis experiments^49^ and by the promiscuous specificity of γ-Thrombin generated by autoproteolysis of α-Thrombin. These cleavages generate an extensive disorder region in the 60-loop which provides an explanation for the observed loss of specificity^50^.

So far, different techniques have been applied for in-depth characterization of the substrate specificity of α-Thrombin, including combinatorial peptide libraries^51–53^ and phage display libraries^54^. In this study, we accurately recapitulated the well characterized α-Thrombin specificity and compared it in the so far most comprehensive fashion the other blood coagulation proteases included in the study. Notably, we increased the knowledge about their substrates/cleavages by a large margin (Table S2, S4), defined their specificities (Figure S4, Table S5), cleavage entropies (Figure S5, Table S6), and block cleavage entropies (Figure S6, Table S7) to show that this unique dataset could recapitulate and extend the knowledge on structural determinants of protease specificity.

### 2.4. Detection of allosteric effects of Na^+^ on the blood cascade proteases

In the previous section we found that specificity profiles generated with our method were sensitive enough to detect subtle differences between the investigated proteases. We next asked whether HTPS could detect changes of specificity profiles corresponding to structural rearrangements of a protease catalytic pocket exemplified by the case of Na^+^-induced allosteric changes in α-Thrombin^55, 56^. Allostery is a crucial regulatory mechanism of proteins where the binding of an allosteric effector modulates conformational and consequently functional changes of a protein. To investigate the effects of Na^+^ on the reorganization of the hydrophobic pocket in the active site of blood coagulation proteases and the ensuing effects on activity and specificity, we generated protease fingerprints of tested coagulation proteases in the presence of 0.2M NaCl or choline chloride (ChCl). ChCl was used to keep the ionic strength constant without exerting an allosteric effect ^57^. Different from other thrombin allostery studies^55^, the reaction was performed at physiological temperature of 37°C and 0.2M NaCl, where about 50% thrombin is bound to Na^+^ molecules^56^, to ensure efficient proteolysis. We first tested whether the experimental conditions recapitulated the well-known activity patterns of α-Thrombin towards Fibrinogen (FGA) and PC. Previous studies have shown that α Thrombin, when bound to Na^+^ (Fast form) has an enhanced activity towards the proteolysis of fibrinogen to fibrin, crucial for clot formation (pro coagulant), but not towards PC, a protease that influences α-Thrombin activity with a negative feedback mechanism^58–60^. As the dissociation constant of α-Thrombin with bound Na^+^ is close to the concentration of the ion in blood at 37°C^58^, a subtle deviation of Na^+^ concentration e.g. around platelet thrombi *in vivo*, generates a different substrate selectivity with an important implication for the pro- vs. anti-coagulatory activity of α-Thrombin. The cleavage patterns observed in our data were used to infer the amino acid specificity enrichment of α-Thrombin towards its known physiological substrates (FGA and PC)^54^. In presence of NaCl we observed lower *p-values* for FGA compared to PC substrate; while in presence of ChCl we identified a similar distribution of *p*-values (Figure 4A). This is in agreement with previous studies which showed that 0.2M Na^+^ led to an increase in the specificity towards FGA, but not towards PC^58^. This together with boosted activity of α-Thrombin in presence of Na^+^ (Figure 4C) results also in a higher rate of FGA cleavages.

**Figure 4.**
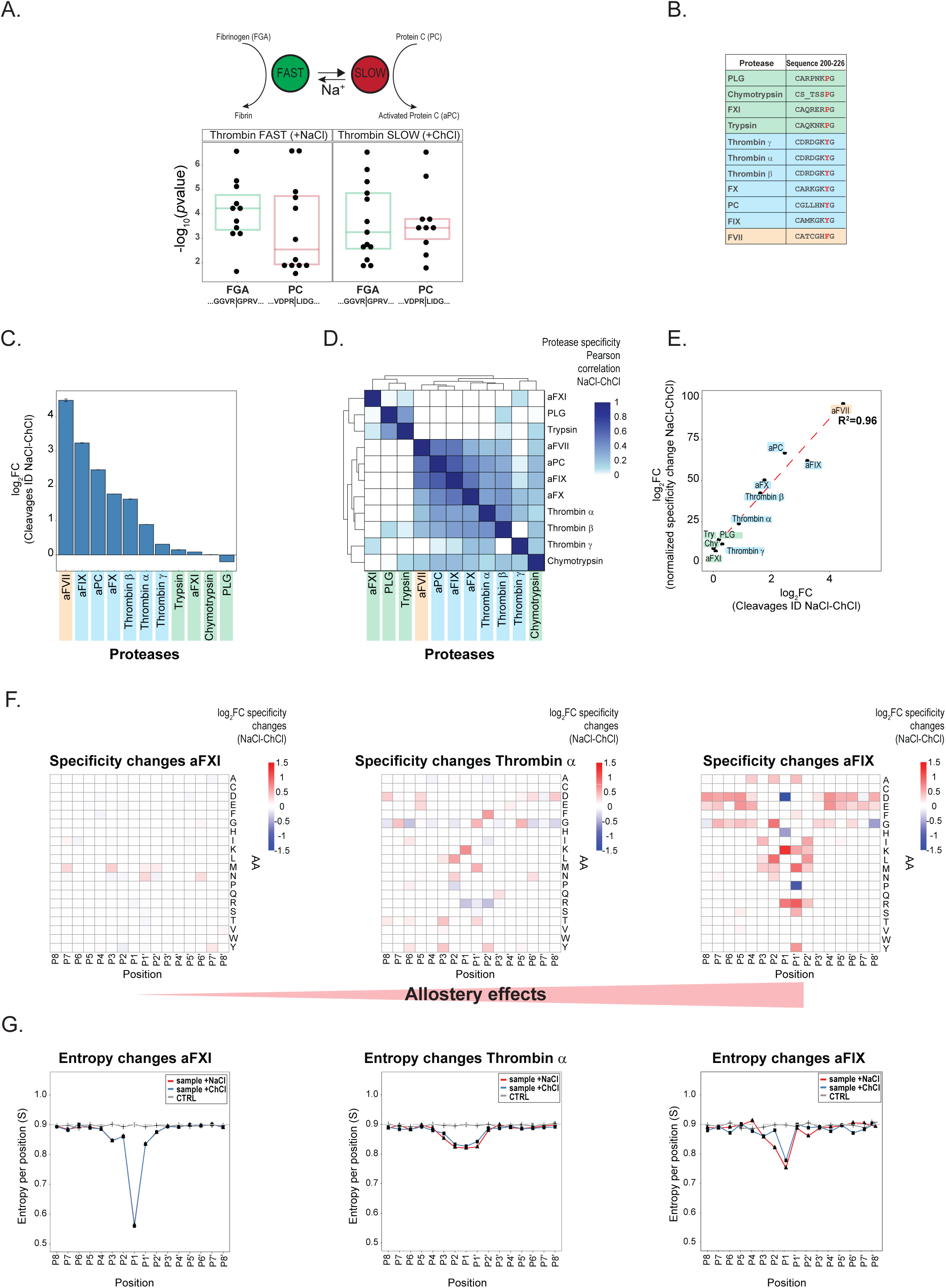
HTPS detects substrate specificity changes induced by Na^+^ allosteric regulation. (A) Effect of Na^+^ in the modulation of α-Thrombin specificity (pro versus anti-coagulant activity) and the equilibrium between the fast and slow forms, respectively. The amino acid specificity enrichment of α Thrombin towards its known physiological substrates (FGA and PC) is characterized by the distribution of *p-values* generated from the positional enrichment of each single amino acid compared to a random distribution. (B) Chymotrypsin like family protease primary sequence alignment in correspondence of the site which regulate the coordination of Na^+^ (220-226), adapted from^61^. Residue 225 is crucial for Na^+^- induced allosteric regulation of serine proteases. Proteases with Tyr (blue) or Phe (orange) in position 225 can coordinate the Na^+^ ion while proteases with Pro (green) in position 225 cannot bind it. (C) Relative log_2_ change of identified peptides in presence and absence of Na^+^. The color code corresponds to the allosteric requirements reported in 4B. (D) Unsupervised hierarchical cluster of protease specificity changes as a result of allosteric effects (NaCl-ChCl). The color code corresponds to the allosteric requirements. (E) Correlation plot between the changes of identified cleavages and changes of substrate specificity observed as a result of Na^+^ allosteric interaction. (F) Heat maps of specific substrate changes generated by Na^+^ allostery: aFXI, activated α-Thrombin and aFIX are ordered based on the magnitude of allosteric effects at the specificity level, where aFXI basically does not show any changes while aFIX shows the maximal changes detected within our dataset. (G) Positional entropy changes generated by Na^+^ allostery: aFXI, activated α-Thrombin and aFIX are ordered based on the magnitude of entropy change (S) observed at the positional entropy level.

Other proteases, similar to α-Thrombin, exhibited an increase in proteolytic activity and these patterns reflected well the requirements for allosteric regulation of blood cascade proteases, where Na^+^ can be coordinated only if Phe or Tyr are at position 225^61–63^ (Figure 4B). Indeed, we observed the strongest fold change of activity (Figure 4C) between NaCl and ChCl in case of aFVII (Phe at position 225), followed by aFIX, aPC, aFX, activated β-, α- and γ-Thrombin (all having Tyr at position 225). In contrast, for Trypsin, aFXI, Chymotrypsin and PLG, proteases with a Pro in position 225, we observed no significant differences between NaCl and ChCl (Figure 4C). Interestingly, we observed that γ-Thrombin, which contains a Tyr in position 225, shows a somewhat intermediate pattern between the two groups, presumably due to the flexibility of the 60-loop which generates a reduced selectivity in the catalytic pocket (Figure 4C, Figure 3A).

We next performed an unsupervised hierarchical clustering to investigate the impact of allostery on the specificity and entropy of blood coagulation proteases included in the study. We clustered the significant specificity changes detected in presence of NaCl vs. ChCl (Figure 4D) and observed that proteases regulated allosterically by Na^+^ clustered closely together and showed significant changes in their substrate specificity. In contrast, no significant changes were observed for proteases that cannot bind Na^+^ and thus cluster separately. The specificity differences observed in case of aFVII, aFIX, aPC, aFX, β- and α-Thrombin suggest, that Na^+^ had an impact not only on the number of cleavages, but also on substrate specificity (Figure S7). Importantly, this is also evident from the correlation between activity changes and the changes detected at the level of substrate preference with a R^2^ value of 0.96 (Figure 4E). Next, we clustered the proteases according to the changes in cleavage entropy in the presence of NaCl or ChCl and found it was in agreement with the allosteric structural constraints of blood coagulation proteases (Figure S8A). Further, we correlated the level of activity change, reflected by the number of detected peptides, with the changes on the level of entropy (Figure S8B). We observed also in this case an R^2^ value of 0.83, demonstrating that proteases where Na^+^ had the most significant impact on activity showed corresponding changes at the level of entropy and specificity. These results convincingly demonstrate that Na^+^ binding to the allosteric site of coagulation cascade proteases regulates their activity in a way that reflects on protease substrate preference (Figure 4F) and entropy (Figure 4G) with strong changes observed, for example, for aFIX, moderate changes for activated α-Thrombin and no changes observed for aFXI (for other proteases, see Figure S5, S6 and S7). The activity and entropy changes showed that in presence of Na^+^ α-Thrombin was more active and had lower entropy values (higher substrate specificity); these data are in line with the mechanism of conformational regulation which suggest an increase in structural rigidity and an open conformation that can stabilize the proteolytically active enzyme^55, 64, 65^.

Overall, we characterized the effect of Na^+^, a critical and well-known allosteric binder and regulator, on all coagulation proteases included in the study, and showed that HTPS has the capacity to detect the functional consequences of allosteric changes at an exquisite level of sensitivity. We found that differential activity, specificity and entropy correlated exactly with the presence/absence of residues that enable Na^+^ coordination. This highlights the potential of this approach as a tool for systematic screening of the effects of drugs or peptide-mimetic molecules as modulators of therapeutically relevant proteases.

### 2.5. Designing fluorescent substrates for blood cascade proteases

To demonstrate the translational value of HTPS we used the protease specificity data generated above to design fluorescent substrates to detect and discriminate the activity of activated α-Thrombin and aFX. The design of fluorescently or otherwise labeled substrates is important to characterize proteases in kinetic cleavage assays and to use this knowledge to support the design of new activity-based probes and inhibitors. After characterizing the specificity of the proteases, we selected for each position amino acids with most significant positional enrichment (Figure 5A, 5B). We designed two synthetic peptides that represent the best match according to the detected specificity for activated α-Thrombin (NH_2_- GIPR↓AAGD-COOH) and aFX (NH_2_-GIGR↓RIAE-COOH). As our analysis investigated the positional specificity but did not take into account possible sub-site cooperativity^53^, we confirmed that these peptides were cleaved effectively by the respective proteases. We monitored the intensity of the cleavage products by mass spectrometry, using MS1 signal integration (Figure 5C). The results showed the expected patterns and thus confirmed that both synthetic peptides represent a good entry point for development of substrates.

**Figure 5.**
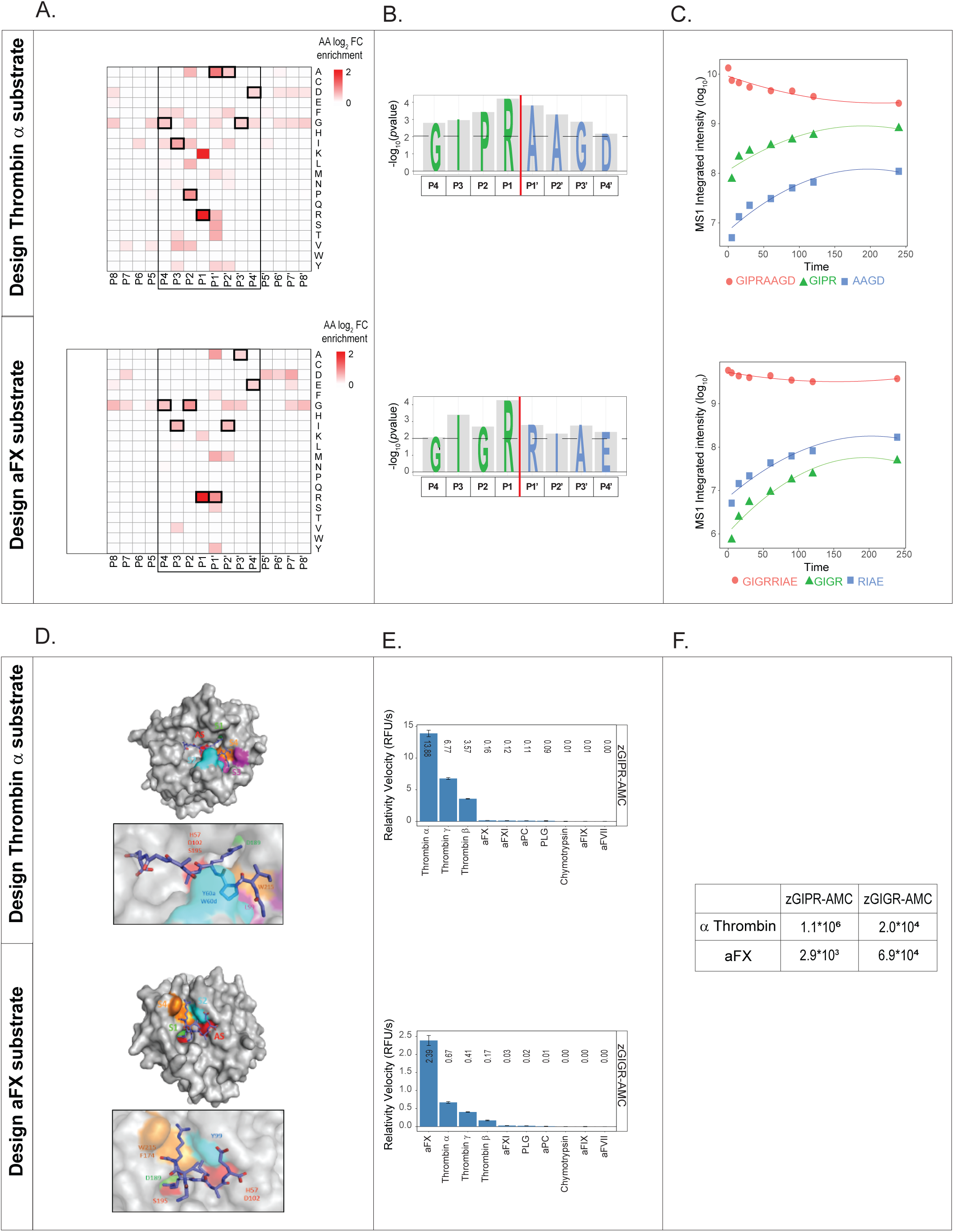
Design of octapeptides and fluorescent substrates. (A) Determination of best-matched substrates from the positional amino acid preferences using positional substrate preferences (log_2_ fold change enrichment compared to random distribution). The best-matched positions selected for the substrate design are highlighted in bold squares. (B) Representation of -log_10_ *p-values* of AA selected for the octapeptide design to assay protease activity of α-Thrombin and aFX. (C) I MS1 intensity integrated area for targeted octapeptide substrates and the corresponding cleavage fragments after incubation with activated α-Thrombin and aFX over time (0 – 240 min). (D) *In silico* docking of activated α-Thrombin and aFX with the two model octapeptides obtained by HPEPDOCK software. The docked peptides are shown in stick mode and proteases (α-Thrombin 1ppb; aFX 1g2l) in surface representation. The location of active site (AS) and specific active pocket sites (S1-S4) are indicated in different colors (green, orange, red and purple). (E) Determination of substrate selectivity measured by the relative reaction velocities (RFU/s) of fluorescent substrate processing for activated α-Thrombin and aFX tested with a panel of closely related coagulation proteases. (F) The calculated k_cat_/K_M_ values for activated α Thrombin and aFX.

To evaluate the exact mode how these peptides bind to the protease active site (AS), we performed a molecular docking analysis. The structural data for α-Thrombin (1ppb) and aFX (1g2l) showed strong similarities of the S1 specific pockets (Asp 189), whereas the S2 and S4 subsites were characterized by distinct topologies. While S2 is covered by the 60-insertion loop in α-Thrombin, it is smaller and solvent accessible in aFX. Further, the aryl-binding site S4 of α-Thrombin, located above the conserved residue Trp 215, is lined by residues Leu 99 and Ile 174. In aFX, the S4 subsite is built by the corresponding residues Tyr 99 and Phe 174, which together with the indole ring system of Trp 215 form the walls of an aromatic box. Our docking models (Figure 5D) showed how the residue P1 (Arg) can be effectively oriented inside the S1 pocket, while the different P2 residues (i.e., Pro in the case of α-Thrombin and Gly in the case of aFX) can fit specifically in the correspondence of S2 subsites, thus ensuring the interaction with the AS. These results indicate that the substrate-design based on HTPS results produces structurally plausible solutions.

Next, we designed small fluorescent tetrapeptide substrates corresponding to the P4-P1 active site preferences to monitor the activity of activated α-Thrombin (zGIPR-AMC) and aFX (zGIGR-AMC). While some selectivity is lost because of placing the fluorophore at P1’-P4’ positions, we tested the substrates against a panel of proteases in a standard assay^66^ and observed that both substrates had selectivity for the target proteases (Figure 5E). Accordingly, zGIPR-AMC was most efficiently cleaved by α-Thrombin and also by γ- and β-Thrombin, while other proteases included in the assay did not cleave zGIPR-AMC. The zGIGR-AMC was less selective because it was cleaved by aFX and also by α-, β- and γ-Thrombin. We further calculated the k_cat_/K_M_ and demonstrated that zGIPR-AMC had good selectivity for α-Thrombin over aFX (380-fold higher k_cat_/K_M_). Selectivity in case of zGIGR-AMC was substantially lower with k_cat_/K_M_ 3.5-fold higher values for aFX in comparison with α-Thrombin (Figure 5F). The measured kinetic parameters of the Thrombin substrate were in the same range as commercially available^67^ and other reported substrates^68^. Accordingly, the α-Thrombin substrate had a 1000-fold higher k_cat_/K_M_ values compared to H< β< AGR< pNA and a 180-fold higher k_cat_/K_M_ value compared to zGGR< AMC, two commercial substrates used in Thrombin generation assays^67^. The chromogenic Thrombin substrate S2238 (H-(D)-Phe-Pip-Arg-pNA), which has physicochemical properties similar to zGIPR-AMC, is 23-fold more selective than our zGIPR-AMC^69^, because it contains in position P3 a non-natural amino acid (D-Phe) that provides additional selectivity in the hydrophobic pocket. This example demonstrated the translational value of HTPS screen, where it is possible to generate a substrate with reasonable selectivity towards the investigated protease in a simple and straightforward way, and without extensive testing or large peptide libraries.

### 2.6. Using HTPS data to predict physiological substrates

For the extensive characterization of the selected target proteases in this study, a full native lysate from HEK293 cells was used. Since coagulation cascade proteases are known to be secreted and their relevant substrates are primarily found in blood, our characterization is likely to have captured a large number of biochemically plausible, but physiologically irrelevant substrates/cleavages. As a final validation step of our protocol, we therefore asked whether we could use the protease specificity information derived from a native lysate to generate hypotheses on proteins known to be secreted into the blood. To pursue this aim, we developed a three-step filtering framework to single out, from a large initial search space, substrate candidates for specific proteases (Figure 6A). We used as a reference the 718 proteins reported as secreted to the blood (human blood secretome from ProteinAtlas^70, 71^) which contain approximately 0.3 million 8- residue sequence combinations (from P4-P4’). In the first step of the analysis, we used a filtering strategy based on the HTPS-generated protease data. Specifically, we calculated positional enrichment of each AA against a random distribution generated from the database and scored each of the potential target sequences using the sum of the significant fold changes associated with the respective residues (Motif score) as shown on Figure 6B. This step correctly identified 13 well-known Thrombin natural targets (PC, FVIII, IGFBP5, FV, FXI, FGA and FGB)^54, 60^ (Figure 6C) in the top 1% of candidate substrates. We observed that the distribution of Motif score was bimodal for promiscuous proteases with high specificity in P1 position (i.e. lower entropy at the cleavage site) such as Trypsin and PLG (Figure S9B, S9C), and much less discrete for those showing less promiscuous specificity features, e.g. α-Thrombin (Figure 6B) and aFVII (Figure S9A) (due to its broad specificity, PLG was excluded from the subsequent analysis). This indicated that for the latter class, there was not a discrete population of preferred substrates, but broader specificities modulated by the combination of all amino acids (cooperativity effects). Using MEROPS (release 12.1) substrates^23^ identified in the secretome as true positives, we constructed receiver-operator curves and filtered the data to match a false positive rate of 1% (Figure 6D). By this means, we could reduce the search space of potential physiological substrates by about 100 times. The evaluation of the sensitivity and specificity indicated a good prediction power for all proteases included in the analysis (average AUC∼0.97, Figure S9D-J). In a second step, we used the JPred4^72^ software tool to predict the solvent accessible regions of (nearly) all 718 secretome proteins and removed all the sequences that were predicted to be buried, thus eliminating structurally implausible targets. This step further reduced the number of potential targets by about a half, from 2688 to 1380 (in case of α-Thrombin substrates). Finally, in a third step, we used loose protein-level filters to further refine the target selection: proteins for which no expression was measured as well as those for which no co-citation with the target protein was reported were removed. A good proxy for physiological substrates was calculated from the ranking of the frequency of co-citation, protein abundance and the number of potential cleavage sites. These steps significantly reduced the search space (we identified for Thrombin 794 potential physiological cleavages), while having a negligible effect on the recall of previously known substrates (Figure S9K). As expected, the final score generated from our filtering strategy was highly skewed towards known substrates reported in MEROPS, indicating that it correctly reports potential substrate candidates (Figure S9L, Table S8). Furthermore, we found that using this filtering strategy, most target sequences were unique to specific proteases and only a few were shared among all six (Figure 6E). Interestingly, the protein substrates displayed an opposite trend: only a few proteins were targeted by a single protease, and the large majority was potentially a target of several or even all of them. The high number of target sequences carried on average by each target protein seems to explain to a significant extent this observation (Figure 6F).

**Figure 6.**
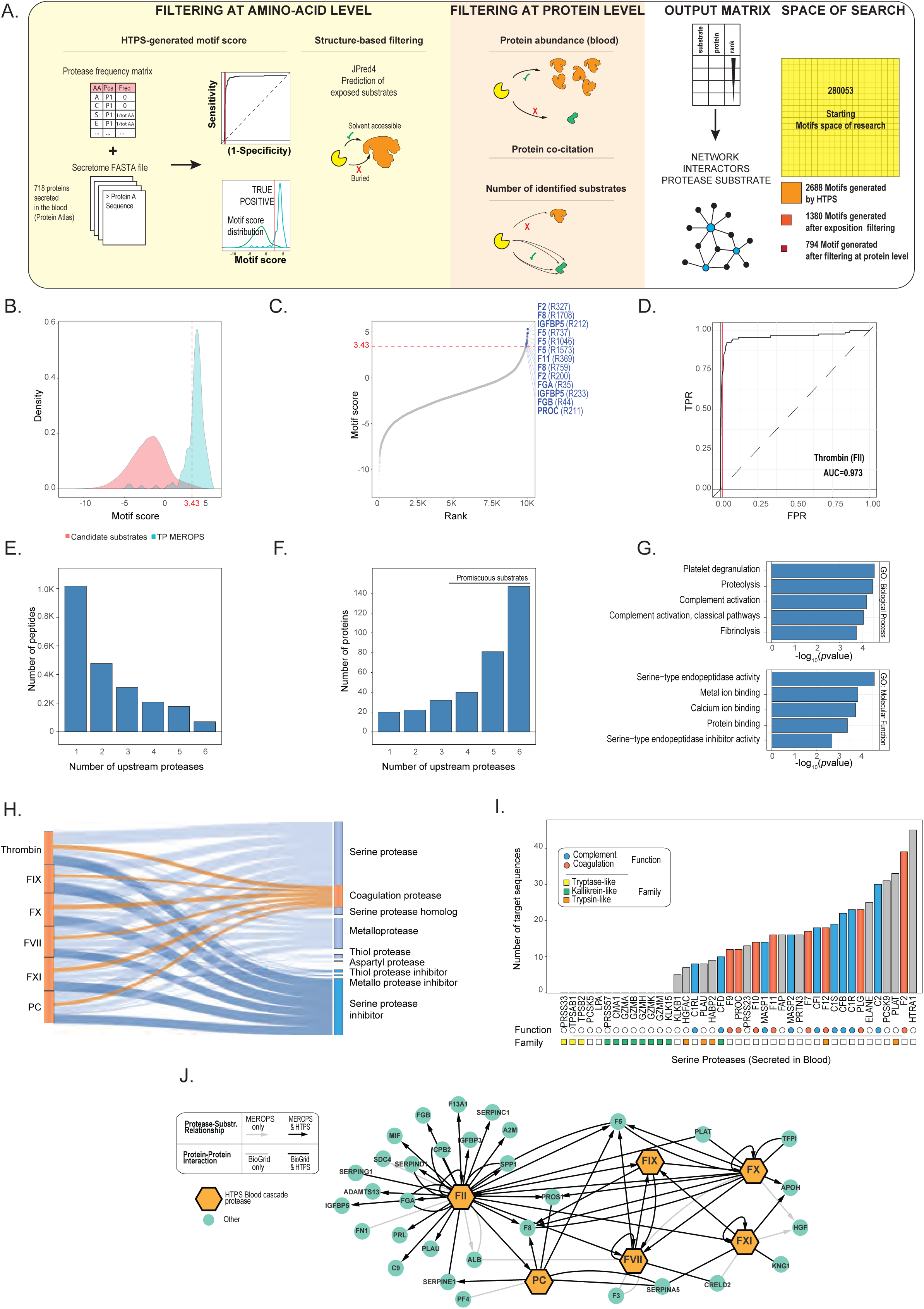
Identification of physiological candidate substrates from HTPS data. (A) The figure shows a three step-filtering framework to identify candidate substrates in the blood secretome. All proteins annotated in ProteomeAtlas as part of the secretome are scored based on the motif score generated by HTPS. The distribution of motif score is filtered using a cut-off of 0.1 FPR using as true positive the cleavages deposited in MEROPS. In the second filtering step, the prediction of amino acid accessibility is used to identify proteases-accessible substrates. In the third step a protein level filter is applied to exclude proteins, for which the concentration in the secretome was not determined (ProteinAtlas database) and/or are not co-cited with the investigated protease. In the final matrix, all proteins are ranked based on the number of identified substrates, protein co-citation and protein abundance. In the right part of the panel, the search space reduction across the three filtering steps (expressed as number of potential substrate sequence for thrombin) is shown. (B) α-Thrombin motif score distribution (light red) and true positive distribution (light blue) calculated from the positional enrichment of each amino acid of all secretome proteins against the motif score generated by HTPS. (C) Distribution of the motif score of α-Thrombin generated for all proteins of the secretome. Annotated α-Thrombin physiological substrates are depicted in blue. The motif score cut-off is depicted by a dashed red line. (D) A Receiver-Operator Curve (ROC) of α-Thrombin to evaluate the performance of the filtering step generated by HTPS motifs. (E) Bar plot of the distribution of the number of peptides identified by the filtering strategy generated by a unique protease or shared between multiple proteases. (F) Bar plot of the distribution of the number of proteins identified by the filtering strategy generated by unique proteases or shared by multiple protease. (G) David GO enrichment of Biological Process and Molecular Function for filtered candidate substrates. (H) Sankey diagram showing the distribution of proteases and protease inhibitor classes across the identified candidate substrates. (I) Bar plot of the number of candidate substrate sequences identified for the 41 serine proteases included in the final filtered list. Proteases associated with complement and coagulation are shown in blue and red, respectively, and main represented families (based on Panther database) are reported as colored squares. (J) A network of coagulation proteases and their identified substrates. The network was generated from protein-protein interaction database (simple connection) (BioGRID v.3.6.1.8.2) and MEROPS substrates database (arrow). Protein substrate information generated from the HTPS data by the previously described filtering steps was superimposed onto the network (black edges).

Next, we asked which processes and functions were enriched among the proteins targeted by the blood cascade proteases. We used DAVID^73^ to calculate the enrichment against the secretome background and found that, in line with our expectations, proteins involved in complement activation and fibrinolysis were enriched among the potential targets, with the class of serine-type proteases being most significantly represented in this subset (Figure 6G). A Sankey diagram (Figure 6H) shows indeed that serine proteases were the main substrates of the investigated blood cascade proteases which displayed generally similar connectivity also with other proteases and protease inhibitor classes. By plotting the number of target sequences for all 41 serine proteases found in blood, a number of additional trends emerged. All proteases that were part of the complement and coagulation pathways were among potential candidates, while about a third of the proteases, especially kallikrein-like and tryptase-like, had no target sequence identified after filtering (Figure 6I). Collectively, the observations about the correctness of scoring and recall of known substrates as well as enrichment of relevant biological processes and functions, indicate that this strategy is able to recover potentially relevant physiological substrates of investigated proteases.

Finally, we combined the knowledge about protease substrate relationships deposited in MEROPS and the information about protein-protein interactions deposited in BioGrid^74^ (v.3.6.1.8.2) in a single network and overlaid it with the data retrieved with our method. Remarkably, as shown in Figure 6J, our analysis was able to capture the large majority of previously known protease-substrate relationships and protein-protein interactions in an entirely data-driven way. We thus defined a strategy to generate context-relevant substrate predictions from HTPS-experimental results obtained in generic systems. This strategy allowed the reduction of the sequence search space by more than 3 orders of magnitude and was able to isolate biochemically, structurally and biologically plausible and thus likely relevant protease-substrates relationships.

## 3. Discussion

Here we describe and benchmark a high-throughput protease screen (HTPS) and demonstrate its performance with selected applications for protease research. We characterized 15 proteases under physiologically relevant conditions and collectively identified more 500.000 substrate peptides and 120.000 unique substrate cleavages, expanding the current knowledge base for all deposited protease studies by a factor of 4. The protocol is simple, scalable, robust, easy to parallelize for multiple conditions (reduces batch effects), avoids any chemical modifications or labeling and, as few biochemical steps are required (no enrichment or depletion), it reduces sample loss. A suite of publicly accessible scripts that support the analysis of the generated data complement the wet lab protocol. As an example, the screening of 10 coagulation proteases in triplicates under three different conditions (without NaCl, with ChCl and with NaCl) required typically 2-5µg of individual tested protease, the native cell lysate of a single 15cm dish (5mg of total proteins) as substrate sample, and could be carried out in only half day of bench work and 2 hours of MS acquisition time per sample. The benchmarking of the method with standard proteomic proteases and metalloproteases has produced two main conclusions. First, HTPS is able to recapitulate accurately protease specificity with similar performance as other methods. Second, HTPS does generally lead to the identification of vastly larger numbers of substrate peptides identified per protease, in comparison with most of the other methods so far used for protease characterization. Furthermore, the highly parallel setup reduces batch effects and increases the method throughput. It can also simultaneously recover prime and non-prime substrate specificity (besides of DIPPS^21^, PICS^75^ and ChaFraTip^22^), but does so in native conditions.

We demonstrate the microscale and high-throughput capabilities of HTPS by applying the workflow to a set of coagulation cascade proteases and detect specificity features for activated α-, β-, γ-Thrombin, aFVII, aFIX, aFX, aFXI, aPC and PLG. Here, the high numbers of detected cleavages allowed us to characterize the minor distinguishing features between these closely related proteases and group them according to their cleavage specificity and cleavage entropy. Furthermore, we were able to recapitulate from our proteomic data the known specificity differences between two isoforms of Thrombin (α- and γ-), which further demonstrated the sensitivity of the screen. The large number of cleavage events identified per measurement allowed us to investigate the allosteric effect of Na^+^ on activity and specificity with great sensitivity: we obtained results that confirm the mechanisms of allosteric regulation for α-Thrombin^53, 58^, aPC^63^ and aFX^62^ and expand our knowledge also to other blood proteases for which so far mechanisms of allosteric regulation with Na^+^ were not extensively described. This demonstrated that differential specificity and entropy profiling can be used to identify restraints to model conformational changes. It is also important to note that allosteric effects are typically investigated with fluorescence anisotropy, biochemical or structural studies which require high amounts of proteases (e.g. in mg range for protein crystallography). In contrast, in its current implementation HTPS analyses are performed in near-native conditions, requires less than 1µg of protease per assay, and further downscaling can be envisioned.

The translational value of HTPS is perhaps best illustrated in the context of designing sensitive tools for detection of protease activity. We used HTPS data to design synthetic peptides and show that they were cleaved by their respective proteases, demonstrating that positional substrate preferences detected with our protocol can translate into tools for detecting protease activity. This is useful, especially in case of poorly characterized proteases where a fast and simple design of a substrate can assist further protease characterization steps. An exemplary application of this concept could be the design of test substrates to characterize proteases of the newly emerging virus. Moreover, large protease datasets could possibly serve as a hypothesis-generator for targeted assays^76^ and for spike-in assays used for detection of protease activity^77^ as recently demonstrated for asparaginyl endopepdidase^78^. Furthermore, protease datasets could support the development of assays that serve as sentinels to follow biological processes in a high-throughput fashion^79^. It must be borne in mind, however, that HTPS is limited to amino acids that naturally occur in proteins in comparison to synthetic peptide libraries. When designing specific substrates for proteases, especially if the target group are closely related proteases, including non-natural amino acids to protease screens is beneficial and can provide another level of selectivity^80^.

As a final, and possibly also the most important application, we show that the large number of identified protease cleavages in near-native conditions can be exploited to predict relevant substrates in systems orthogonal to those experimentally used. Here, a simple computational filtering framework, largely based on HTPS-results, combined with readily available orthogonal information was capable to retrieve a large number of physiologically relevant relationships and generate hypotheses on yet unexplored connections. The substantial pool of substrates/cleavage events identified with HTPS may play in the mid-term also an important role in bringing machine learning approaches to protease research and improve the performance of tools readily used for prediction of protease substrates. Recent developments of tools like iProt-Sub, that can predict cleavages in protein substrates, demonstrated the importance of having a detailed and representative cleavage dataset for the investigated proteases to retrieve their specificity features and thus construct better models that could enable proteome-wide prediction of protease substrates^81^.

To conclude, we introduce a new proteomic tool for protease research, which we dub HTPS. We believe it could be readily applied for large-scale de-orphaning of proteases, systematic comparison of their specificity and entropy, identification of potential physiological substrates, as well as generation of substrate reporters to investigate protease activity and structural rearrangements. Further improvements, including adoption of more sensitive MS, shorter LC gradients and scaling down of the starting material, will make profiling of the entire human protease repertoire across different conditions a goal within reach, as only ∼9 sets of experiments would, in principle, be sufficient to profile it in triplicates on a 384-well format.

## 4. Materials and methods

### Proteases used in the study

All proteases used in this study were purchased from commercial vendors, Trypsin (V5111), Glu-C (V1651), Asp-N (V1621) and Chymotrypsin (V1091) from Promega (USA) and Lys-C (125-05061) from Wako (Japan). The blood cascade proteases α-Thrombin (HCT-0020), β-Thrombin (HCBT-0022), γ-Thrombin (HCGT-0021), Factor VIIa (HCVIIA-0031), IXa (HCIXA-0050), Xa (HCXA-0060), Xia (HCXIA-0160), Plasmin (HCPM-0140) and activated Protein C (HCAPC-0080) were purchased from Hematologic Technologies, Inc., (USA). Recombinant human MMP-2 (902-MP-010) was purchased from R&D systems (USA). Recombinant human MMP-3 (SRP7783) was purchased from Sigma Aldrich (Germany).

### Cell culture and preparation of native cell lysates

Human Embryonic Kidney 293 cells (HEK293, ATCC CRL-1573) were grown under standard conditions in DMEM (Gibco) supplemented with 10% FBS (BioConcept), 1% glutamine (Gibco), and 1% penicillin/streptomycin (Gibco) at 37°C in a humid incubator at 5% CO_2_. When the cells reached 90 % confluence, they were detached from the plate with a jet of PBS (Gibco) and washed twice with PBS. For lysis, we used mild lysis conditions with HNN buffer (50 mM HEPES, 150 mM NaCl, 50 mM NaF, pH 7.8) supplemented with 0.5 % NP-40 and protease inhibitor cocktail according to manufacturer’s recommendations (Sigma Aldrich) as described elsewhere^82^. Afterwards, the lysate was centrifuged at 14.000g for 15 minutes to remove any non-soluble material and the buffer was exchanged for 20 mM Ammonium Bicarbonate pH 7.8 using a filter device with molecular weight cutoff of 10 kDa. Standard BCA protein assay was used to determine the total protein concentration (Thermo Fischer Scientific), the concentration of the standardized lysate was set to 1 mg/ml and stored at −80 °C until used for the digestion assays.

### Protease digestions and sample preparation

All protease digestions were performed in 96FASP plates with MWCO 10 kDa (Acroprep Advance^TM^) by adapting a 96FASP sample preparation protocol for protease digestion under native conditions^28, 29^. The first step was to wash the filter units to remove any residual glycerol. For this 100 µl of 20 mM Ammonium bicarbonate pH 7.8 were added to the wells and the plate was centrifuged at 1300g for 10 minutes before repeating this step once more. Afterwards, native cell lysate standardized in 20 mM Ammonium bicarbonate pH 7.8 was added at a final 50 µg of total protein per well and mixed with the investigated proteases at 1/50 [E]/[S] ratio. The samples were incubated at 37 °C for 4 hours before the flow-through was collected by a 15 min centrifugation at 1300g in a low binding 96-well conical plate. The collection step was repeated by adding 100 µl of MS-grade water to the wells for collecting the flow- through. The fractions were transferred to low-binding tubes (Eppendorf) and concentrated on the SpeedVac to complete dryness. The samples were stored at −80 °C until analysis. Before analysis, the samples were re-suspended in 20 µl of MS-grade water with 0.1 % formic acid and the peptide concentration was determined with Nanodrop UV spectrometer. The sample concentration was adjusted to 1 µg/µl with water with 0.1 % formic acid.

### Allosteric effects of Na^+^ on blood cascade proteases

To investigate the effect of Na^+^ on the proteases of the blood cascade we performed the assay in presence of 0.2M of NaCl or choline chloride (ChCl) as previously reported^55^. We performed the digestion experiments with blood cascade proteases under both conditions for 2 hours at 37 °C in 96-well plates and collected the peptides as previously described. Additionally, before the LC-MS/MS analysis we performed a desalting step of the samples with C18 UltraMicroSpin columns according to the manufacturer’s protocol (The Nest group, USA). The dried peptide samples were re-suspended in 0.1% FA water at a concentration of approximately 1µg/µL.

### LC-MS/MS analysis

The LC-MS/MS analysis of the protease-digested samples was performed on an Orbitrap Elite (Thermo Fischer Scientific) interfaced with an Easy 1000 nano-LC unit (Thermo Fischer Scientific), coupled online with the nano-electrospray. The LC-MS/MS was operated with the Xcalibur software package (Thermo Fischer Scientific). For the analysis, 1 µg of sample was loaded directly on the analytical column (Acclaim PepMap^TM^ RSLC, 75 µm x 15 cm, nanoViper C18, 2µm, 100A, Thermo Fischer Scientific). The flow rate on the nano-LC was set to 300 nl/min and the peptides were chromatographically separated with a 5 – 35 % 120 minute linear acetonitrile/water gradient in 0.1% formic acid. During the entire run, the MS spectra were acquired in the Orbitrap in positive ion mode with 2.0 kV voltage in the mass range of 350 to 1600 m/z, set to the profile mode and a resolution of 120.000 at 400 m/z. For peptide fragmentation, a CID fragmentation method with normalized collision energy 35 was used and the MS/MS spectra were obtained from the 15 most intense precursor ions from the full MS spectra. During the entire run, precursors with repeat count of 1 were dynamically excluded for 30 seconds. Precursors with charges +2, +3 and +4 were considered and the MS/MS spectra were recorded in the ion trap analyzer in the centroid mode with normal scan rate and standard settings.

### Database searches and abundance-focused library generation

The raw data was searched with MaxQuant^31^ (version 1.5.2.8) using the human UniprotKB database (*Homo sapiens*, Uniprot release 2018_10, 20382 entries) and the in-house generated abundance-focused *HTPS_DB.fasta* database (2557 entries). For specific database searches we used standard MaxQuant settings^83^, the searches without a defined enzyme specificity were performed as described elsewhere^21^. Our searches considered only two natural PTMs, acetylation of N-termini (+42.0106 Da) and the oxidation of methionine (+15.9949 Da) as variable modifications. First search peptide mass tolerance was 20 ppm and main search peptide mass tolerance was 4.5 ppm, as set by default. MS/MS match tolerance was set to 0.5 Da. For the peptide identification via peptide-spectrum matching the FDR was controlled with a standard target-decoy approach^83^. A 1 % peptide FDR was applied at PSM level and only peptide hits with a PEP score ≤ 0.05 and a score > 40 were retained for further analysis. Potential contaminants were not included in the analysis.

For generation of the *HTPS_DB.fasta* abundance-focused database, we combined the lists of proteins that were identified in the samples after treatment with Trypsin, Lys-C, Asp-N, Glu-C and Chymotrypsin. The final list of was the union of proteins identified in the respective samples and we included only proteins with a global protein PEP ≤ 0.01 into the final database.

### Data analysis and visualization

Data analysis was performed in R (version 3.4.3) using the workflow deposited on Github (https://github.com/anfoss/HTPS_workflow) under MIT license. Briefly, the script recovers the cleavage sequences from the identified peptides and transfers them into a positional matrix (amino acids upstream the cleavage site occupy P8-P1 position and amino acids downstream P1’-P8’ position). A frequency matrix is generated counting the abundance of amino acids per position and normalized for all identified peptides. To harmonize the multivariate protease specificity data, the positional occurrences of amino acids are converted into protease frequency matrices. In parallel, a random frequency matrix of the same size is generated by sampling the same number of amino acids as contained in the frequency matrix from the natural distribution of amino acids in *HTPS_DB.fasta*. The proteases were first compared in terms of numbers of generated cleavages under different tested conditions. For visualization of the specificity, we used the iceLogo program^38^ with the threshold of significance *p*-value set to 0.01, respectively. To compare proteases in terms of significantly different positional features, a two-side paired t-test was employed to evaluate similarity between frequency matrices or differential frequency matrices and thus to evaluate the similarities/differences between the tested proteases. The evaluation and comparison of substrate specificity for MMP2 and MMP3 with PICS, TAILS and DIPPS data was performed by adapting the workflow used for HTPS. Identified cleavages or peptides reported in the studies^21, 41–43^ were used to generate the frequency matrix and the specificity enrichment using as a control the random distribution of amino acids from *HTPS_DB.fasta* database. For conditional protease comparison, we took the significant (*p-value* < 0.01) enrichment of amino acid per position compared to the random distribution in presence of NaCl and ChCl, compared the folds of change and report the significant changes according to *p*-value. The calculation of cleavage entropy was performed as a Shannon entropy calculation^34^. The block entropy calculation was performed as described elsewhere^35^.

### Spike-in octapeptides and fluorescent substrates for α-Thrombin and Factor X

The octapeptides GIPRAAGD (α Thrombin) and GIGRRIAE (Factor Xa) were synthesized by the solid-phase method using the 9-fluorenylmethyloxycarbonyl (Fmoc) strategy on a model PS3 automated synthesizer from Protein Technologies International (Tucson, AZ), according to a standard protocol described elsewhere^84^. The crude peptides were subsequently purified by RP-HPLC on a C18 analytical column (Grace-Vydac, Hesperia, CA) and analyzed by MS with a DDA approach. In order to determine the linear response range for the proteases, the two peptide were tested from 100 µM to 10 pM. To confirm the octapeptide cleavage we incubated 10 µM of the peptide with with 10 nM final concentration of proteases from 0 – 240 minutes. For the analysis, 1 µg of sample was loaded directly on reverse phase column (75 µm x 15 cm, packed with Magic C18 3 µm resin) and the peptides were separated with a 5 – 35 % 20 minute linear acetonitrile/water gradient in 0.1% formic acid with a flow rate set to 300nl/ml, using a Proxeon EASY< nLC II chromatography system (Thermo Fischer Scientific). The acquisition started with sample injection. The MS1 quantification of selected reporters was performed on an Orbitrap XL (Thermo Fischer Scientific) in positive mode with 2.0 kV voltage in the mass range of 150 to 1200 m/z in the profile mode at a resolution of 60.000 at 400 m/z. The measurement was performed using 1 µl of the standardized sample spiked with iRT peptides (Biognosys AG) at 1:20 and proteolyzed BSA 0.1mg/ml as carrier. We manually integrated the precursor isotope peaks (M, M+1, M+2) using Skyline software^85^ of GIPRAAGD (378.70 m/z), GIPR (221.64 m/z), AAGD (333.14 m/z) for α Thrombin and GIGRRIAE (436.26 m/z), GIGR (201.63 m/z), RIAE (488.28 m/z) for aF10.

The fluorescent substrates for α-Thrombin (zGIPR-AMC) and for aFX (zGIGR-AMC) were purchased from Biomatik (USA) and selectivity was tested in a standard protease screen as described elsewhere^66^. All measurements were performed in 20 mM Ammonium bicarbonate pH 7.8 with 200 mM NaCl. Where applicable, we also determined the corresponding k_cat_/K_M_. The substrate concentration range in the assays was 1 µM - 200 µM, the protease concentration range was 1 - 5 nM. We monitored the increase of fluorescence intensity with a Tecan infinite 2000 Pro plate reader (Tecan, Switzerland) in continuous mode (excitation at 370 nm, emission at 460 nm) and calculated the corresponding k_cat_/K_M_ values as earlier described^66^.

### *In silico* data analysis

Molecular docking was performed with HPEPDOCK web server^86^, starting from the structures with the water molecules removed and inhibitor-free for α-Thrombin (1ppb)^87^ and aFX (1g2l)^88^ and the two octapeptides. The software generated 3D structure models for the given sequences of peptides using the implemented MOPEP program, which considers peptide flexibility. Simulations were run with default parameters, without introducing any geometric or energetic constraints. One hundred poses were generated and ranked according to the CAPRI criteria^89^. The most acceptable prediction was selected for the data analysis. PyMOL software (v. 0.99rc6) was used for visualization of the docking results.

### Identification of candidate substrates

To identify physiologically relevant protease substrates we applied three filtering steps. In the first filtering step, we calculated a motif score for all the combination of amino acids (280053) of all secretome proteins (secretome database from Protein Atlas^70, 71^). The motif score for each protease analyzed (Trypsin, Thrombin, aFVII, aFIX, aFX, aFXI, PLG and aPC) was calculated from the sum of significant fold changes associated with the respective residues compared to a random distribution generated from HTPS database. To evaluate the performance and to identify a cut-off at 1% FPR we generated a receiver operator curve (ROC) using as true positive the annotated MEROPS substrates (release 12.1)^23^. The identified substrates were further filtered based on the prediction of amino acids exposition using JPred4 tool^72^ (http://www.compbio.dundee.ac.uk/jpred/). For this filter step, we used the intermediate score “JNETSOL_5” for the accessibility prediction and we filtered all substrates that were not buried (*n*=8). In the last step, we applied a protein-based filter. In this step we removed proteins for which the concentration in blood was not reported (ProteinAtlas, Secretome^70, 71^, https://www.proteinatlas.org/) and/or were not co-cited with the studied individual coagulation protease in PubMed (https://www.ncbi.nlm.nih.gov, ftp://ftp.ncbi.nlm.nih.gov/gene/DATA/gene2pubmed.gz). Furthermore, proteins were scored by multiplying the inverse of ranking position for i) co-citation frequency, ii) number of identified protease substrates, iii) concentration in the blood. GO enrichment for Biological Process and for Molecular Function was performed using DAVID tool^73^ (v6.8, https://david.ncifcrf.gov/) using the human secretome (ProteinAtlas) as background. Protease substrate network was generated using Cytoscape (v.3.8.0)^90, 91^, combining data of reported protein interaction in BioGRID (v.3.6.1.8.2)^74^ and substrates MEROPS (v.12.1)^23^ for all coagulation protease.

## Abbreviations

AA: Amino acid
AMBIC: Ammonium bicarbonate
AS: Active site
ChaFraDIC: Charge-based fractional diagonal chromatography
ChaFraTip: ChaFraDIC performed in a pipet tip format
COFRADIC: Combined fractional diagonal chromatography
DDA: Data Dependent Acquisition
DIPPS: Direct in-gel profiling of protease specificity
FASP: Filter-aided sample preparation
96FASP: FASP performed in a 96-well plate format
FC: Fold change
FDR: False discovery rate
FPPS: Fast profiling of protease specificity
HEK293: Human Embryonic Kidney 293 cells
HTPS: High-throughput protease screen
MMP: Matrix metalloprotease
MQ: MaxQuant
MWCO: Molecular weight cut-off
PICS: Proteomic identification of protease cleavage sites
PSM: Peptide-spectrum match
TAILS: Terminal amine isotopic labeling of substrates

## Acknowledgements

R.A. and M.V. are supported by the Swiss National Science Foundation through grant # SNSF 31003A_166435 to R.A. and by the European Research Council (ERC) through grant 20140AdG 670821. M.G., F.F. and F.U. acknowledge support by the IMI project ULTRA-DD (FP07/2007-2013, grant no. 115766). U. adK. acknowledges support by a Novo Nordisk Foundation Young Investigator Award (NNF16OC0020670). This work was also supported by a Grant from the CaRiPaRo Foundation Excellence Research Project 2018 BPiTA n. 52012 to V.D.F.

## Author contributions

M.V., F.U. and R.A. conceived the study. M.V. and F.U. performed the experiments, F.U., M.V, R.C. and A.F analyzed the data. A.F. and F.F. contributed to the experimental design. L.A. and V.D.F. synthesized the substrate peptides and provided valuable input for experiments. U. adK. provided information for MMPs and valuable discussions. M.V. and F.U. wrote and R.A. revised the manuscript. M.G. provided funding. All authors edited the manuscript. R.A. supervised the project and provided funding.

## Data Availability

The data is deposited to ProteomeXchange with the identifiers PXD018976 and PXD020320.

## Code availability

The R scripts to analyze the data and reproduce the reported data analysis are available at https://github.com/anfoss/HTPS_workflow under MIT license.

