## Supplementary Figures for "Mapping specificity, entropy, allosteric changes and substrates in blood proteases by a high-throughput protease screen"

#### SUPPLEMENTAL FIGURES

**Figure S1. Quality control of [Glu-1]-Fibrinopeptide B human (GF).**

Quality control of GF peptide external standard. The sensitivity of the MS instrument and column performances were checked using the intensity (A) and the retention time (B) of GF peptide.

**Figure S2. Target library generation for HTPS and evaluation of performance for specific and unspecific database searches.** (A) The target library (HTPS_DB) size focused on the abundant proteome covering around 12% of the human Uniprot *Homo sapiens* database (2557 proteins). (B) Distribution of residues abundance in Uniprot database and in HTPS_DB database. (C) Distribution of peptide FDR for identified peptide using unspecific searching mode in MaxQuant for Uniprot database (blue line) and HTPS_DB (yellow line). The analysis was performed for a sample generated by trypsin (upper panel) and chymotrypsin proteolysis (lower panel). (D) Numbers of identified peptides (FDR < 0.01) for unspecific and specific searches using Uniprot database and HTPS_DB database. The analysis was performed for samples generated by trypsin (upper panel) and chymotrypsin (lower panel) proteolysis.

**Figure S3. Influence of original peptide termini on specificity profiles**. Correlation of the fold of change enrichment of detected substrate preference for (A) Trypsin, (B) Chymotrypsin, (C) MMP2 and (D) MMP3 generated from all detected peptide termini (FC_HTPS_peptide) and from cleavage sequences (FC_HTPS_cleavage).

**Figure S4. Heat maps of positional substrate preferences of blood cascade proteases.** The heatmaps show the $log_2$ fold of change in preference/enrichment for amino acid at investigated positions (P8-P8') in comparison to a random distribution. The respective natural amino acids are sorted alphabetically.

**Figure S5. Positional entropy profile plots of studied proteases.** The plots show positional entropy values (S) for each position (P8-P8') for condition without NaCl (green), with NaCl (red), with ChCl (blue) or for a control sample (grey) for all proteases included in the HTPS screen calculated according to Fuchs et al.[34]

**Figure S6. Block entropy profile plots of studied proteases.** The plots show block entropy values for each position (P8-P8') for condition without NaCl (green), with NaCl (red), with ChCl (blue) or for a control sample (grey) for all proteases included in the HTPS screen calculated according to Qi et al.[35]

**Figure S7. Specificity protease change generated by $Na^+$ allostery.** The positional specificity change of NaCl vs. ChCl is shown for positions P8-P8' for all coagulation proteases included in the screen. The differences are depicted as the $\log_2$ FC for each amino acid for each substrate position for the tested conditions.

**Figure S8. HTPS detects substrate specificity changes induced by $Na^+$ allosteric regulation as reflected at the entropy level.** (A) Unsupervised hierarchical cluster of protease cleavage entropy changes as a result of $Na^+$ allosteric regulation (NaCl-ChCl). The colors correspond to the allosteric requirements: Proteases with Tyr (blue) or Phe (orange) in position 225 bind sodium while Pro (green) in position 225 prevents sodium binding. (B) Correlation plot between the changes detected on the level of identified cleavages and changes detected on the level of substrate entropy observed as the result of $Na^+$ allosteric interaction.

**Figure S9. Evaluation of the performance of three filtering steps to identify potential physiological substrates.** (A) aFVII (B), Trypsin and (C) PLG motif distribution (red) and true positive distribution (light blue) calculated from the positional enrichment of each amino acid of all secretome proteins against the motif score generated by HTPS. (D-J) Receiver-operator curves (ROC) to evaluate the performance of the first filtering step using Motif score generated from HTPS-identified substrate preferences. (K) Summary table with statistical features (precision and recall) of different filtering steps applied at the level of amino acids or at the level of proteins. (L) Distribution of protein rank for known (true positive) substrates annotated in MEROPS (light blue) compared to all identified substrates (light red).

### Figure S1

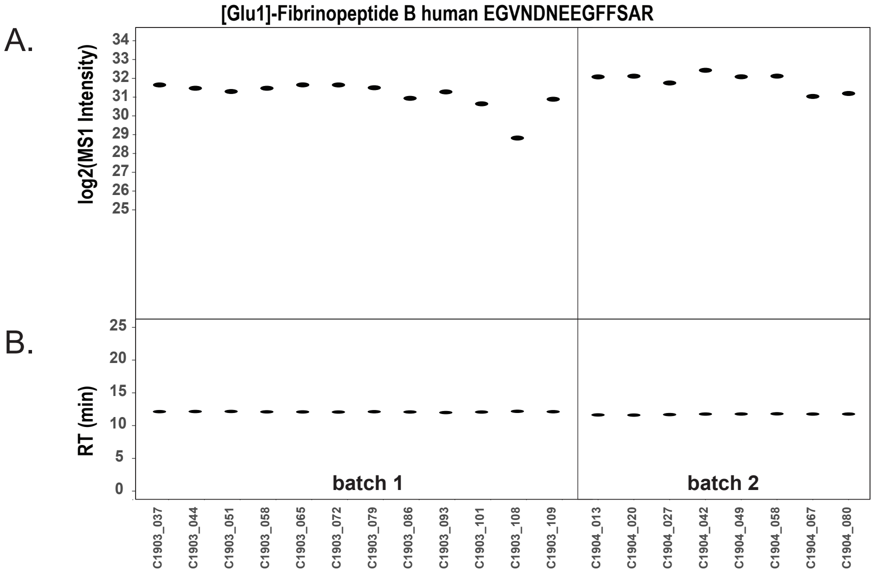


### Figure S2

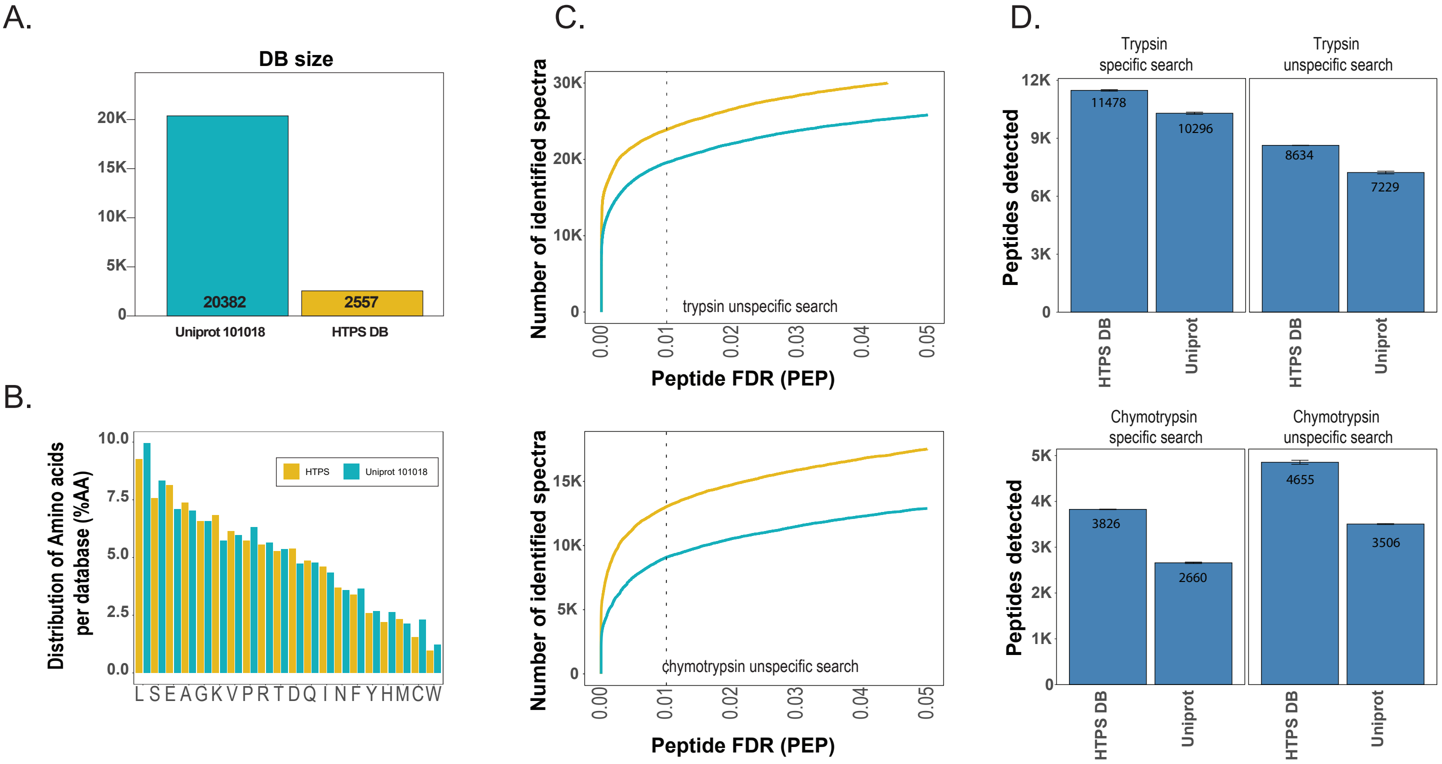

#### Figure S3

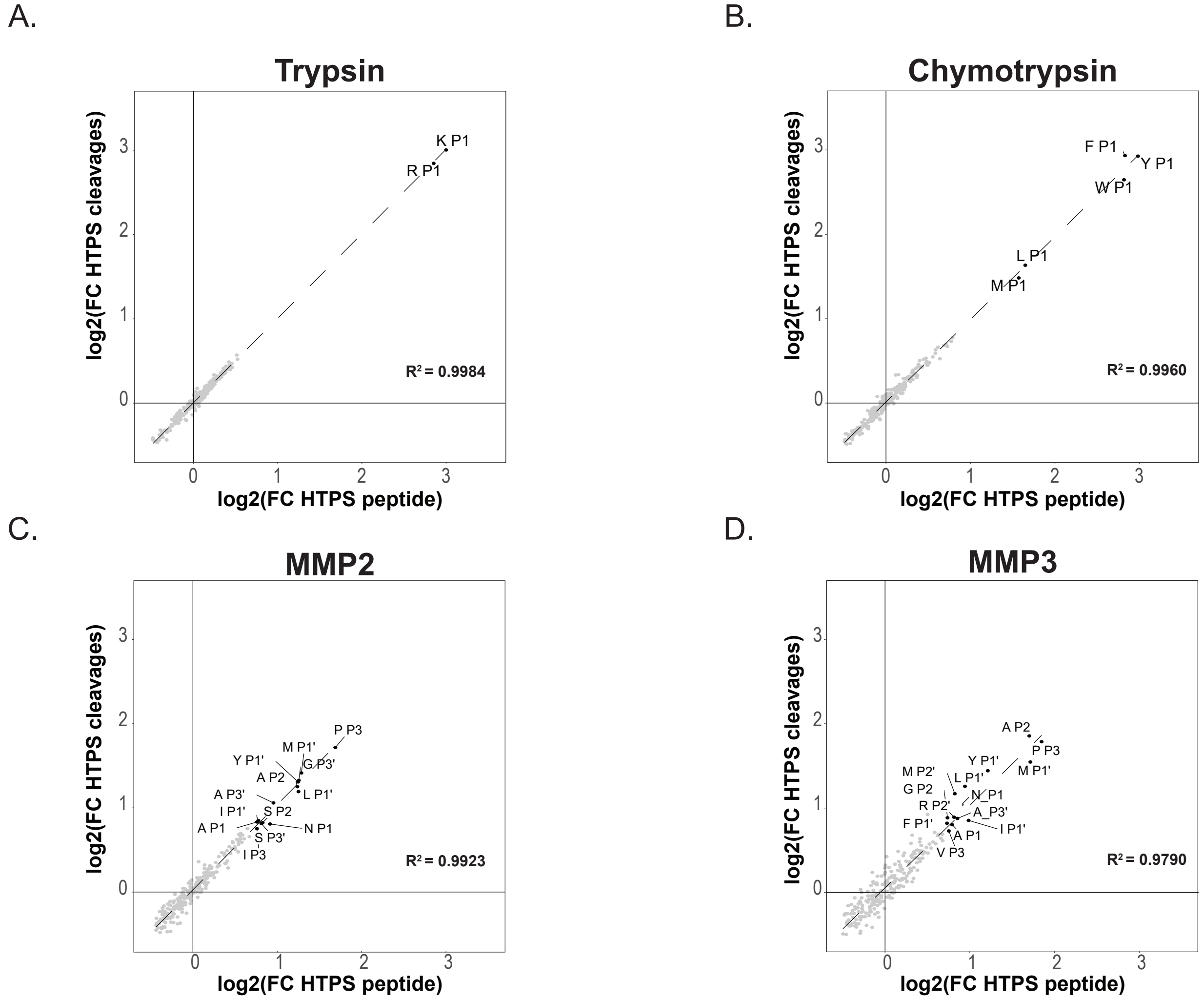
A.
Trypsin
K P1
R P1
R² = 0.9984
B.
Chymotrypsin
F P1
Y P1
W P1
L P1
M P1
R² = 0.9960
C.
MMP2
P P3
M P1'
Y P1'
G P3'
A P2
L P1'
A P3'
I P1'
S P2
A P1
N P1
S P3'
I P3
R² = 0.9923
D.
MMP3
A P2
P P3
Y P1'
M P1'
M P2'
L P1'
G P2
N P1
R P2'
A P3'
F P1'
I P1'
A P1
V P3
R² = 0.9790
log2(FC HTPS cleavages)
log2(FC HTPS peptide)

### Figure S4

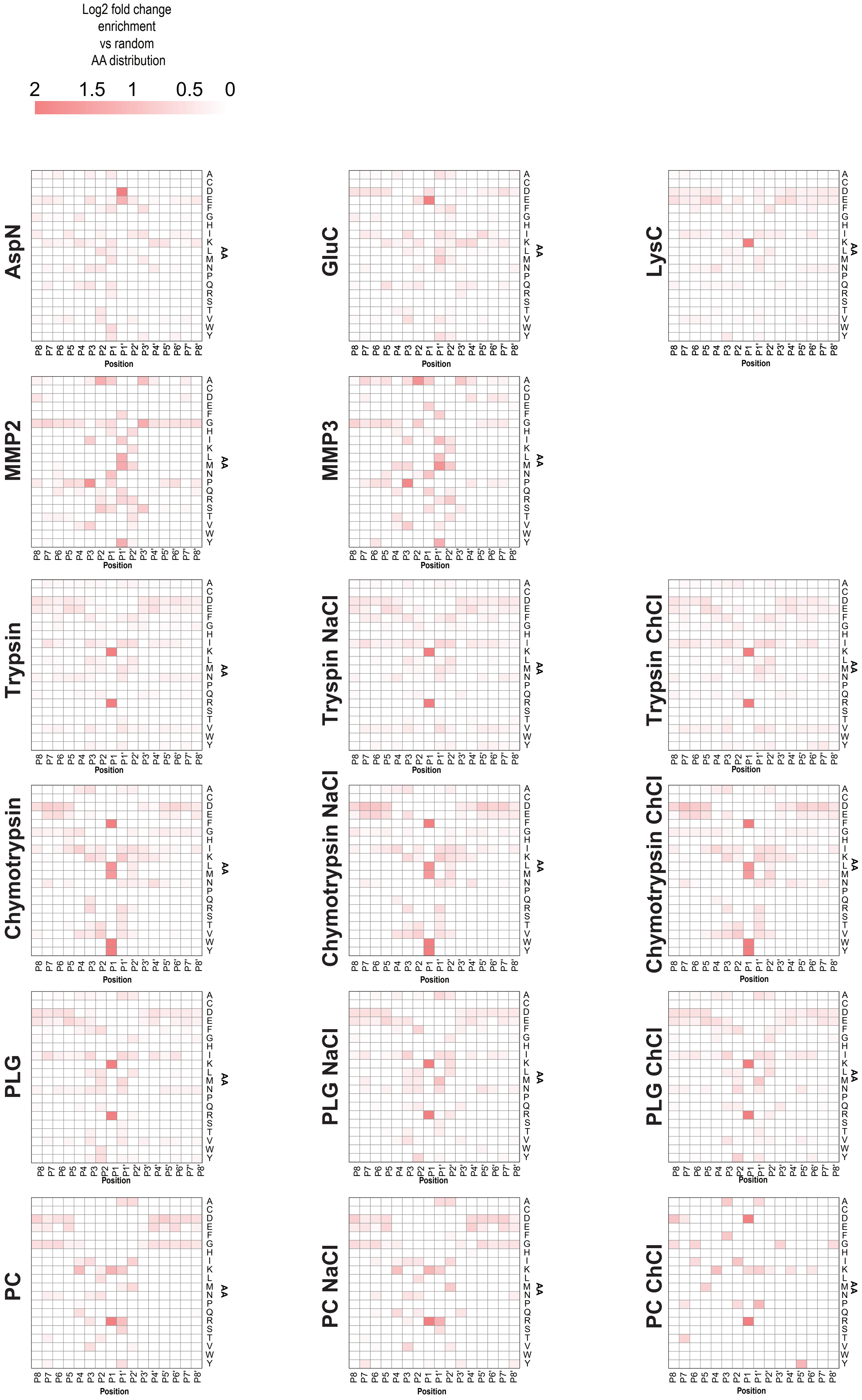

Log2 fold change
enrichment
vs random
AA distribution
2
1.5
1
0.5
0
AspN
GluC
LysC
MMP2
MMP3
Trypsin
Tryspin NaCl
Trypsin ChCl
Chymotrypsin
Chymotrypsin NaCl
Chymotrypsin ChCl
PLG
PLG NaCl
PLG ChCl
PC
PC NaCl
PC ChCl
P8 P7 P6 P5 P4 P3 P2 P1 P1' P2' P3' P4' P5' P6' P7' P8'
Position
A C D E F G H I K L M N P Q R S T V W Y
AA

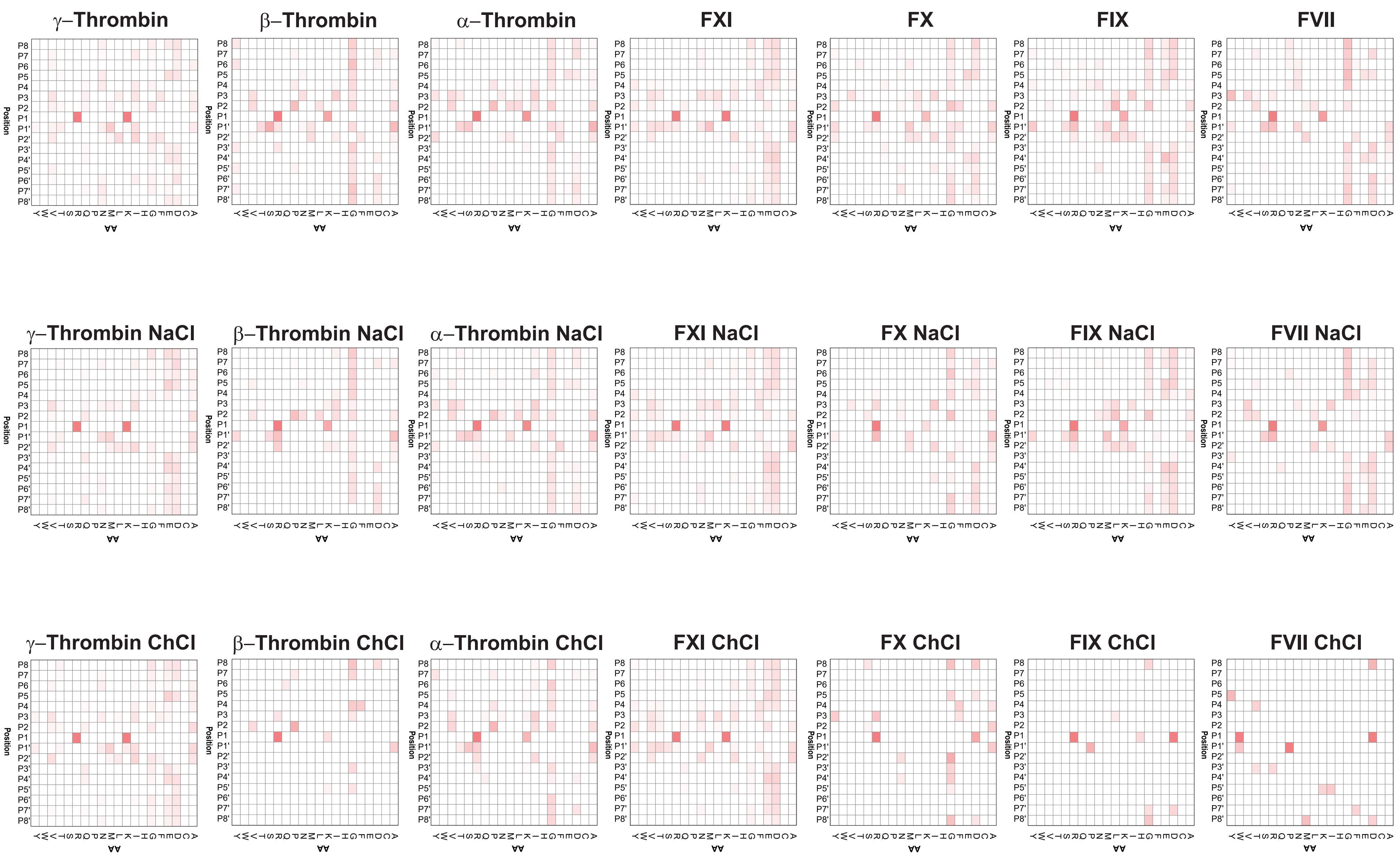
γ–Thrombin
β–Thrombin
α–Thrombin
FXI
FX
FIX
FVII
γ–Thrombin NaCl
β–Thrombin NaCl
α–Thrombin NaCl
FXI NaCl
FX NaCl
FIX NaCl
FVII NaCl
γ–Thrombin ChCl
β–Thrombin ChCl
α–Thrombin ChCl
FXI ChCl
FX ChCl
FIX ChCl
FVII ChCl
Position
AA

### Figure S5

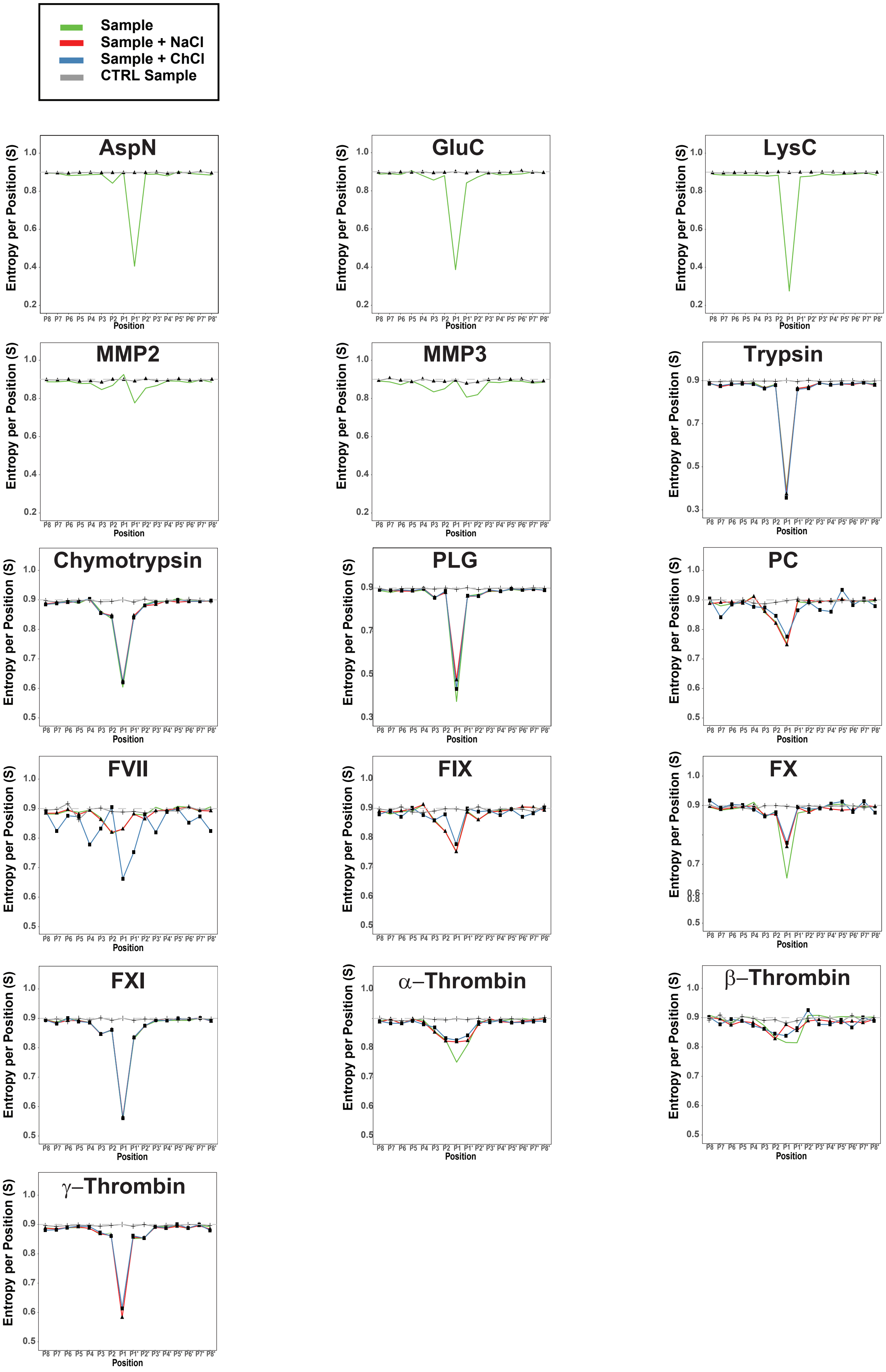

Sample
Sample + NaCl
Sample + ChCl
CTRL Sample
AspN
GluC
LysC
MMP2
MMP3
Trypsin
Chymotrypsin
PLG
PC
FVII
FIX
FX
FXI
α−Thrombin
β−Thrombin
γ−Thrombin
Entropy per Position (S)
Position
P8 P7 P6 P5 P4 P3 P2 P1 P1' P2' P3' P4' P5' P6' P7' P8'

### Figure S6

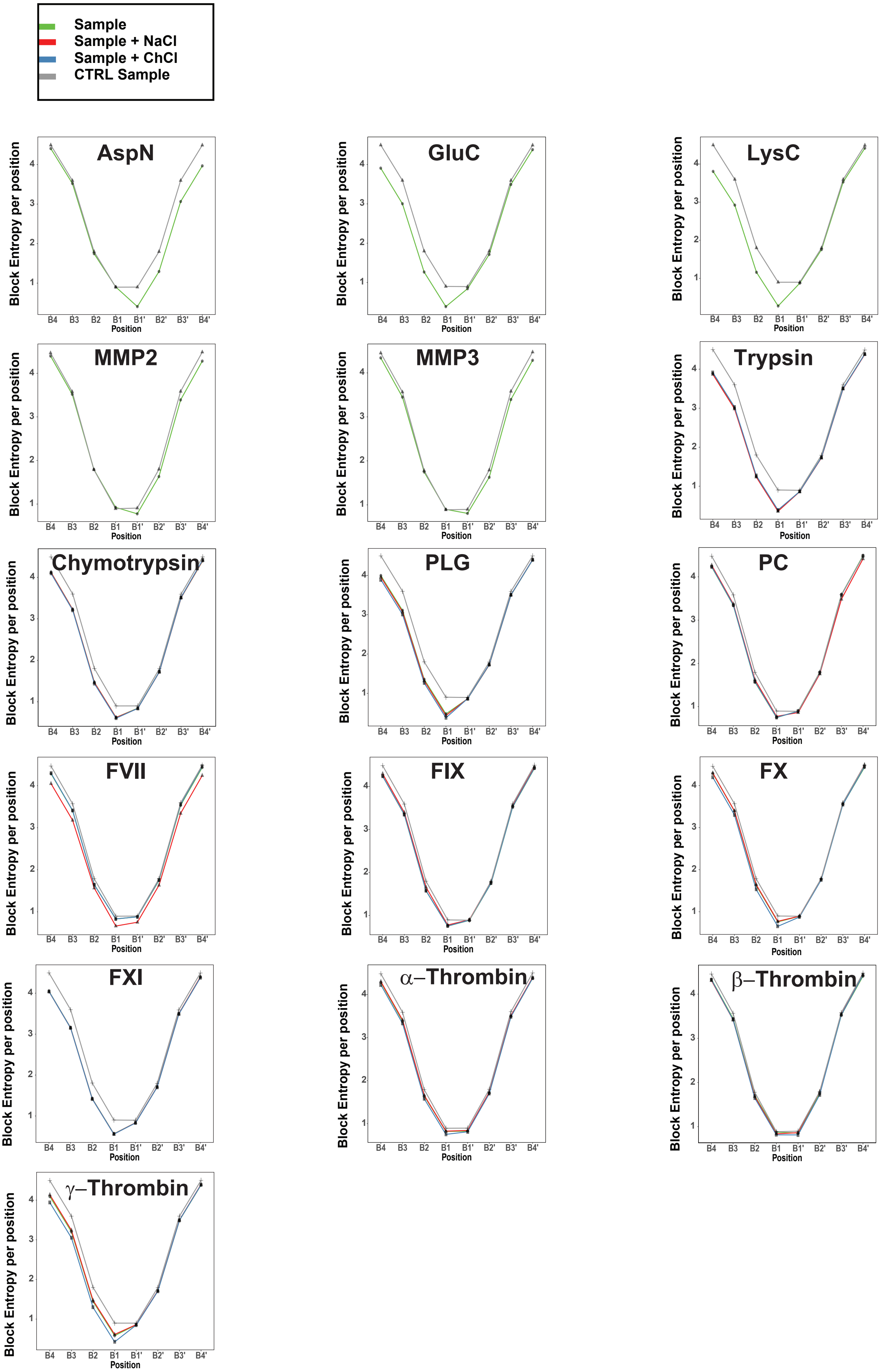

Sample
Sample + NaCl
Sample + ChCl
CTRL Sample
AspN
GluC
LysC
MMP2
MMP3
Trypsin
Chymotrypsin
PLG
PC
FVII
FIX
FX
FXI
α–Thrombin
β–Thrombin
γ–Thrombin
Block Entropy per position
Position
B4 B3 B2 B1 B1' B2' B3' B4'

### Figure S7

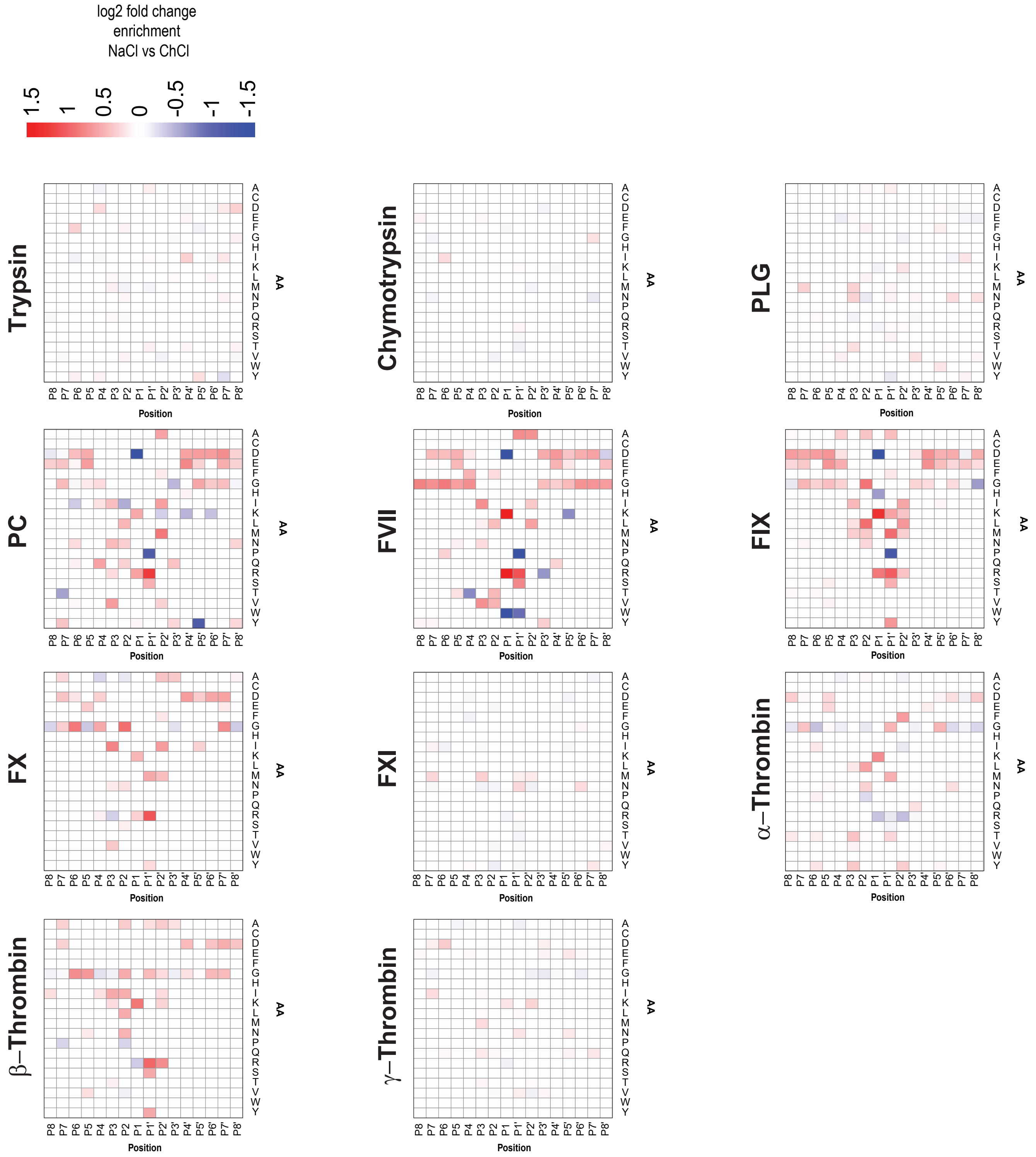


### Figure S8

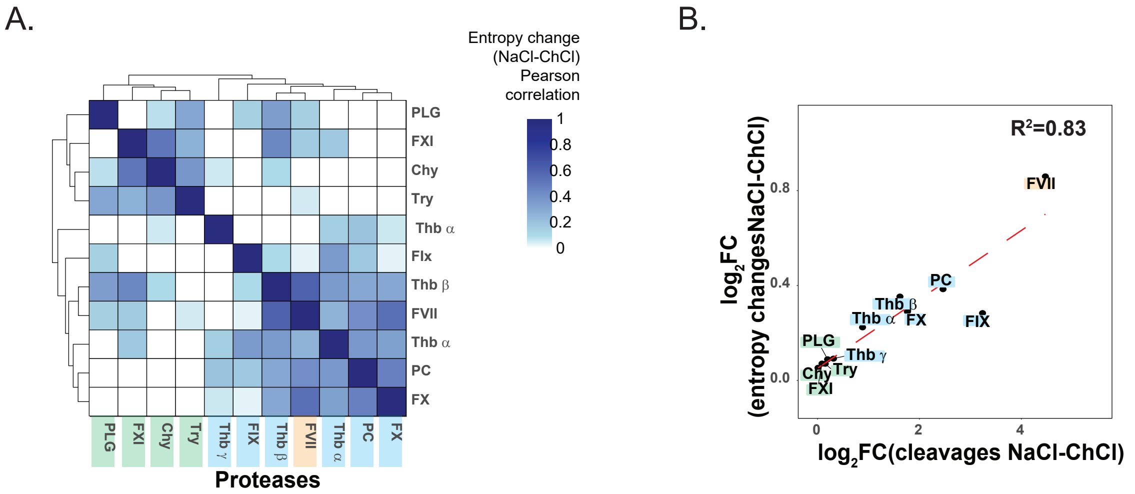

### Figure S9

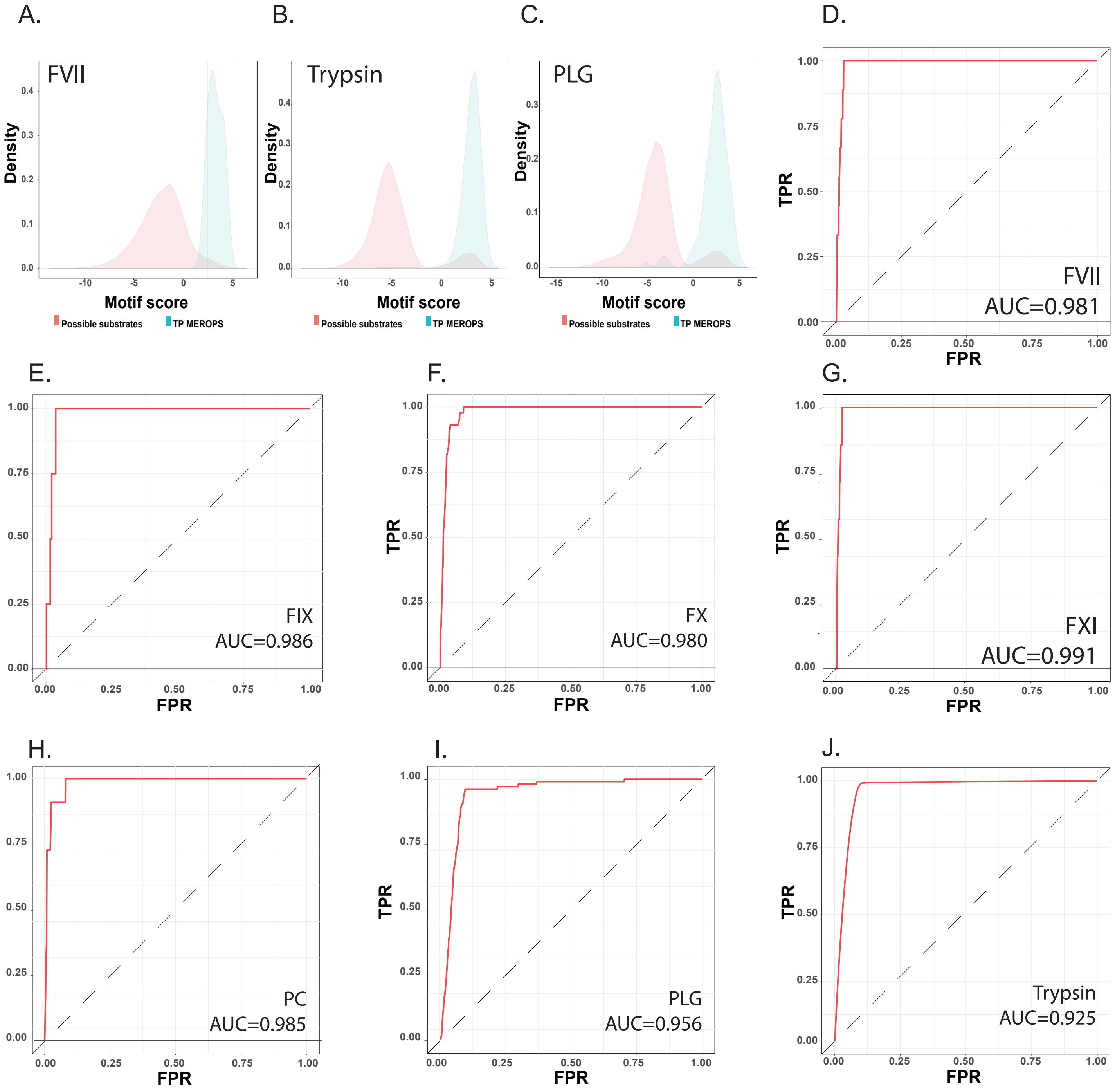


K.

| | | After amino acid level filter | | | | | After protein level filter | | | |
|---|---|---|---|---|---|---|---|---|---|---|
| Protease | TP MEROPS | Threshold (FPR) | Motif score @ threshold | ID motifs | Precision (%) | Recall (%) | ID motifs | ID proteins | Precision (%) | Recall (%) |
| Thb | 41 | 0.01 | 3.433 | 1380 | 1.88 | 63.41 | 794 | 265 | 2.90 | 56.10 |
| FVII | 8 | 0.01 | 3.426 | 1382 | 0.14 | 25.00 | 825 | 267 | 0.24 | 25.00 |
| FIX | 3 | 0.01 | 3.743 | 1383 | 0.07 | 33.33 | 839 | 276 | 0.12 | 33.33 |
| FX | 17 | 0.01 | 3.761 | 1473 | 0.20 | 17.65 | 869 | 269 | 0.23 | 11.76 |
| FXI | 6 | 0.01 | 3.696 | 1249 | 0.16 | 33.33 | 794 | 265 | 0.25 | 33.33 |
| PC | 8 | 0.01 | 3.474 | 1547 | 0.32 | 62.50 | 911 | 265 | 0.55 | 62.50 |
| PLG | 69 | 0.05 | 2.413 | 6697 | 0.60 | 57.97 | 4390 | 354 | 0.84 | 53.62 |
| **average (no PLG)** | **14** | **0.01** | **3.589** | **1402** | **0.46** | **39.20** | **839** | **268** | **0.71** | **37.00** |

L.

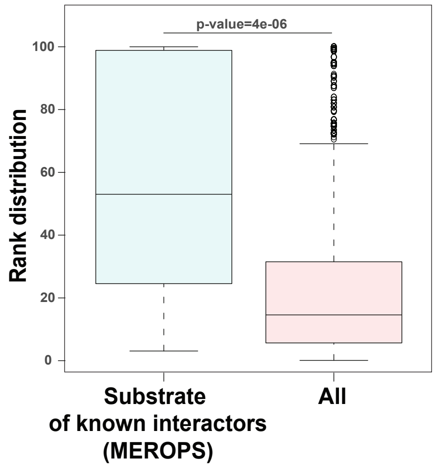
